# Decreased GABA levels during development result in increased connectivity in the larval zebrafish tectum

**DOI:** 10.1101/2024.09.11.612511

**Authors:** Yang Liu, Yongkai Chen, Carly R. Duffy, Ariel J VanLeuven, John Branson Byers, Hannah C. Schriever, Rebecca E. Ball, Jessica M. Carpenter, Chelsea E. Gunderson, Nikolay M. Filipov, Ping Ma, Peter A. Kner, James D. Lauderdale

## Abstract

γ-aminobutyric acid (GABA) is an abundant neurotransmitter that plays multiple roles in the vertebrate central nervous system (CNS). In the early developing CNS, GABAergic signaling acts to depolarize cells. It mediates several aspects of neural development, including cell proliferation, neuronal migration, neurite growth, and synapse formation, as well as the development of critical periods. Later in CNS development, GABAergic signaling acts in an inhibitory manner when it becomes the predominant inhibitory neurotransmitter in the brain. This behavior switch occurs due to changes in chloride/cation transporter expression. Abnormalities of GABAergic signaling appear to underlie several human neurological conditions, including seizure disorders. However, the impact of reduced GABAergic signaling on brain development has been challenging to study in mammals. Here we take advantage of zebrafish and light sheet imaging to assess the impact of reduced GABAergic signaling on the functional circuitry in the larval zebrafish optic tectum. Zebrafish have three *gad* genes: two *gad1* paralogs known as *gad1a* and *gad1b*, and *gad2.* The *gad1b* and *gad2* genes are expressed in the developing optic tectum. Null mutations in *gad1b* significantly reduce GABA levels in the brain and increase electrophysiological activity in the optic tectum. Fast light sheet imaging of genetically encoded calcium indicator (GCaMP)-expressing *gab1b* null larval zebrafish revealed patterns of neural activity that were different than either gad1b-normal larvae or *gad1b*-normal larvae acutely exposed to pentylenetetrazole (PTZ). These results demonstrate that reduced GABAergic signaling during development increases functional connectivity and concomitantly hyper-synchronization of neuronal networks.

**Significance Statement:** Understanding the impact of reduced GABAergic signaling on vertebrate brain development and function will help elucidate the etiology of seizure initiation and propagation and other neurological disorders due to the altered formation of neural circuits. Here, we used fast light sheet imaging of larval zebrafish that neuronally expressed a genetically encoded calcium indicator (GCaMP) to assess the impact of reduced GABA levels through null mutation of *gad1b* during brain development. We show that reduced GABA levels during development result in increased functional connectivity in the brain.

## Introduction

In the functionally mature brain, GABAergic interneurons gate information flow and mediate network dynamics in a variety of contexts, including the processing of sensory information (Yokoi et al., 1995; Isaacson and Strowbridge, 1998; Flores-Herr et al., 2001; Schoppa and Urban, 2003; Lee and Zhou, 2006; Arevian et al., 2008; Popova, 2015), motor control (Beck and Hallett, 2011), and cognition (Tepper et al., 2008; Tremblay et al., 2016; Koyama and Pujala, 2018; Swanson and Maffei, 2019). In humans, reduction in GABAergic signaling is implicated in several pathologies of the central nervous system, including altered sensory processing (Vucinic et al., 2006; Arevian et al., 2008; Puts et al., 2015), aberrant motor control (Solimena et al., 1990; Kim et al., 1994; Levy et al., 1999; Lynex et al., 2004; Beck and Hallett, 2011; Puts et al., 2015), seizure disorders (Ben-Ari, 2006; Glykys et al., 2009; Galanopoulou, 2010; de Curtis and Avoli, 2016; Wang et al., 2017), Tourette syndrome (Puts et al., 2015), autism spectrum disorder (Abrahams and Geschwind, 2008; Geschwind, 2009; Gaetz et al., 2014; Robertson et al., 2016), and schizophrenia (Wu and Sun, 2015). Despite the impact on human health, the effects of reduced GABAergic signaling on the development and function of inhibitory circuits in a live brain are poorly understood.

Genetics offers an unbiased approach to investigating neural function; however, it has been challenging to establish a model system to study the developmental and physiological effects associated with the genetic reduction of GABA synthesis. In vertebrates, GABA is synthesized from glutamic acid by the enzyme glutamic acid decarboxylase (GAD, IUBMB Enzyme Nomenclature EC 4.1.1.15) (Erlander et al., 1991). In mammals and most vertebrates, the majority of GAD protein exists as two molecularly distinct forms, known as GAD67 and GAD65, encoded by the *GAD1* and *GAD2* genes, respectively (Legay et al., 1986; Erlander and Tobin, 1991). In humans, homozygous mutations in the *GAD1* gene are associated with seizures and hypertonia, presumably due to reduced synaptic GABA (Lynex et al., 2004; Chatron et al., 2020); however, high-resolution cellular imaging and developmental studies are not feasible in people. No mutations have been reported for the human *GAD2* gene. Although mice are an excellent genetic model for studying mammalian neural development and function, the role of the *Gad1* gene has been difficult to study as animals homozygous null for *Gad1* die at birth due to cleft palate (Asada et al., 1997; Condie et al., 1997), likely caused by central nervous system (CNS) dysfunction (Oh et al., 2010). While transgenic and conditional workarounds have been developed to study *Gad1* gene function in the pancreas (Yoon et al., 1999), these approaches have not been widely used to perturb GABA synthesis in the brain. Recently, a conditional approach has been used to effect global knockdown of GAD67 in adult mice, which resulted in increased motor activity and impairment of acoustic startle responses as assessed by behavioral assays (Miyata et al., 2021). Mice homozygous mutant for *Gad2* are viable, maintain normal levels of GAD67 and GABA in their brains, and exhibit normal general behavior, including locomotor activity (Asada et al., 1996).

Here we use larval zebrafish homozygous null for *gad1b* and calcium imaging to assess the impact of reduced GABAergic signaling on the function of intrinsic circuits in the optic tectum of the larval zebrafish. The optic tectum of the larval zebrafish is well suited for experiments investigating the functional behavior of circuits. The zebrafish larval tectum integrates and processes visual information for export to premotor targets (Nevin et al., 2010). The tectum is accessible to electrophysiological recordings, and the entirety of the tectum can be imaged at cellular resolution for many hours in the live, intact, non-anesthetized larva (Niell and Smith, 2005; Del Bene et al., 2010; Tao et al., 2011; Gabriel et al., 2012; Nikolaou et al., 2012; Hunter et al., 2013; Muto et al., 2013; Naumann et al., 2016; Vanwalleghem et al., 2018; Burgstaller et al., 2019; Kramer et al., 2019; Liu et al., 2019b; Forster et al., 2020; Wu et al., 2020), which permits assessment of the dynamics of large neuronal populations in response to different challenges. Owing to work by several labs over the last 30 years, including efforts to generate zebrafish brain atlases (Ronneberger et al., 2012; Randlett et al., 2015; Marquart et al., 2017; Kunst et al., 2019a), much is known about cell type diversity and functional connectivity in the zebrafish optic tectum (Nevin et al., 2010; Thompson et al., 2016; Hildebrand et al., 2017; Helmbrecht et al., 2018; Kunst et al., 2019a), which facilitates cell type identification in imaging data. Additionally, recent work has established that spontaneous activity in the optic tectum of the zebrafish larva reveals significant features of the functional connectivity of different circuits (Marachlian et al., 2018). Although the patterns of spontaneous activity are similar to those of visually evoked responses and are organized according to the tectum’s retinotopic map, the formation of the basic circuits does not require visual input or intrinsic retinal activity (Niell and Smith, 2005; Ramdya and Engert, 2008; Grama and Engert, 2012; Avitan et al., 2017; Pietri et al., 2017).

Unlike mice and humans, zebrafish have three gad genes that encode glutamic acid decarboxylase. In addition to *gad2*, zebrafish have two copies of the *gad1* gene, which are *gad1a* and *gad1b* (Grone and Maruska, 2016; Lai et al., 2016; Lai et al., 2017). While it is known that gad1b is expressed by neurons in the larval zebrafish optic tectum (Higashijima et al., 2004; Yu et al., 2011; Barker and Baier, 2015; Forster et al., 2017), *gad1a* expression has not been assessed.

In this study, we test the hypothesis that *gad1b*-null mutations result in a localized expansion of the activity in tectal micro-circuits involving *gad1b*-expressing neurons. We imaged *gad1b*-null mutant larvae expressing a calcium reporter with a light-sheet microscope and quantified the activity level and connectivity between different regions by measuring the correlations in activity. We then compared the connectivity to wild-type larvae and wild-type larvae treated with PTZ. We see an altered pattern of activity in the optic tectum of the *gad1b*-null mutants. Compared to wild-type larvae, the *gad1b*-null mutants show increased connectivity between regions on the same side of the brain and regions on opposite sides.

## Materials and Methods

### Genetic nomenclature

Specific references to genes for humans, mice and zebrafish follow gene nomenclature conventions appropriate for each organism (Mullins, 1995; Bult et al., 2019; Bruford et al., 2020). Human gene symbols are in upper-case italicized characters. Mouse gene symbols are italicized, with only the first letter in upper-case. Zebrafish gene symbols are in lower-case italicized characters. Protein symbols for human and mouse are denoted by upper-case letters not italicized. Protein symbols for zebrafish are not italicized, and the first letter is in upper-case.

### Zebrafish care and maintenance

Adult and larval zebrafish (*Danio rerio*) were obtained from lines maintained in the University of Georgia Zebrafish Facility following standard procedures (Westerfield, 2007). Embryos and larvae were staged using standard staging criteria (Kimmel et al., 1995; Westerfield, 2007). Wild-type fish of the WIK strain and *nacre(mitf)*^w2/w2^ were originally obtained from the Zebrafish International Research Center (ZIRC). Crystal zebrafish (*nacre^w2/w2^*, *alb^b4/b4^*, *roy^a9/a9^*) (Antinucci and Hindges, 2016) were obtained from Dr. Hindges. Fish mutant for *scn1lab* (Baraban et al., 2013; Grone et al., 2017) were obtained from Dr. Scott Baraban. Fish transgenic for TgBAC[*gad1b*: loxP-DsRed-loxP-GFP] (Satou et al., 2013) were obtained from Dr.Shin-ichi Higashijima. Fish transgenic for Tg[*elavl3:GCaMP5g*] (Ahrens et al., 2013a; Ahrens et al., 2013b) were obtained from Dr. Ahrens. Zebrafish mutant for *gad1a* or *gad1b* were generated as previously described (VanLeuven et al., 2018; O’Connor et al., 2019). All adult fish were maintained in an Aquatic Habitats (Apopka, FL) multi-rack system. Habitat water consisted of reverse osmosis filtered/sterilized water to which sodium bicarbonate and other salts (Instant Ocean, Aquarium Systems, Inc., Mentor, OH, USA) were added to maintain pH from 7.0-7.4 and conductivity between 400 and 430 µS. All experimental procedures were conducted in accordance with National Institutes of Health guidelines for use of zebrafish in research under protocols approved and overseen by the University of Georgia Institutional Animal Care and Use Committee.

### Genotypes used for calcium imaging

Since pigmentation interferes with calcium imaging, pigmentation in the larvae used for imaging was reduced using both genetic and pharmacological means. *nacre(mitf)*^w2/w2^, Tg[*elavl3:GCaMP5g*] larvae were used as controls for experiments in which neural activity was perturbed using PTZ. We were not able to generate a line of fish in which the *gad1b* mutant allele was on a nacre background. Therefore, it was necessary to treat *gad1b^−/−^*; Tg[*elavl3:GCaMP5g*] larvae with 0.003% PTU in egg water starting at 18 hpf to suppress pigmentation. The solution was changed once daily until 5dpf. Fish were moved back into egg water before imaging. As a control for possible effects of *nacre(mitf)*^w2/w2^, larvae harboring Tg[*elavl3:GCaMP5g*] but otherwise wild-type were imaged after exposure to PTU as outlined above. No significant differences in calcium activity were observed between *mitf^+/+^*, Tg[*elavl3:GCaMP5g*] larvae reared in 0.003% PTU and *nacre(mitf)*^w2/w2^, Tg[*elavl3:GCaMP5g*] larvae either in the absence or presence of PTZ.

### Colorimetric *in situ* hybridization

Whole mount and section mRNA *in situ* hybridizations were performed as previously described for zebrafish larvae treated with 0.003% PTU (Thisse and Thisse, 2008). Section in situs included an antigen retrieval step added as described in James et al. (2016). Color development was done using NBT/BCIP substrate.

### HPLC-ECD sample preparation

Adult and 7 dpf larval zebrafish were anesthetized in 0.4% Tricaine-S (MS 222; tricaine; pH 7.4) (Westerfield, 1993) and then placed on a pre-chilled metal block. For larval samples, single heads were removed, rinsed with 40 μL of Hank’s Final solution (Westerfield, 1993) and then placed in a pre-weighed 1.5 mL microcentrifuge tube to record the wet mass in milligrams (mg). For adult samples, the heads were removed and brains were dissected out with forceps and rinsed with ~40 μL of Hank’s Final solution. Adult brains were briefly blotted on a piece of filter paper and then placed in a pre-weighed 1.5 mL microcentrifuge tube to record the wet mass in mg. For these preparations, we either added 200 μL of 0.2 N perchloric acid to detect catecholamine neurotransmitters or 200 μL of 18.2 Ω Milli-Q Water to detect amino acid neurotransmitters. Once the solution is added to the tube and samples are fully immersed into the solution, the tubes were immediately frozen on dry ice and stored at −80°C until they were run in HPLC with electrochemical detection (HPLC-ECD). Samples were normalized and run as described previously (Ross and Filipov, 2006; Coban and Filipov, 2007).

### High Performance Liquid Chromatography with Electrochemical Detection (HPLC-ECD)

Concentrations of brain amino acids were determined using high performance liquid chromatography with electrochemical detection (HPLC-ECD; Waters Alliance equipment e2695 and 2465, Milford, MA). Brains were removed, homogenized in 200 ml of MilliQ water, and centrifuged (13,200 x G at 4° C for 10 min) prior to sample supernatant collection. Sample supernatants were made electrochemically active with a derivatizing agent 10 min before sample injection (20 ml) into the HPLC-ECD for detection of glutamine, glutamate, and GABA (Monge-Acuña and Fornaguera-Trías, 2009). The analytes were separated on a C_18_, 5 µm base deactivated reverse-phase column (4.6 µm × 250 mm; Xterra Shield RP18, Waters) using an isocratic flow rate of 0.5 mL/min. The mobile phase with a final pH of 4.5 (adjusted with 1 M phosphoric acid) consisted of 0.1 M monosodium phosphate and 0.5 mM EDTA with 25% methanol (v/v) water (Monge-Acuña and Fornaguera-Trías, 2009). Prior to statistical analysis, amino acid levels were normalized to tissue weight.

### PTZ dose response assay

For assays with wild-type, *gad1b*^+/−^, *gad1b*^−/−^, *gad1a^−/−^*, we either performed a *gad1b* heterozygous incross and post-hoc genotyped each fish from each per treatment group or we crossed several zebrafish of known genotype (wild-type, *gad1b*^−/−^ or *gad1b*^−/−^ x wild-type). Each experimental replicate was performed on separate occasions. For assays with *ga2404* +/− and *ga2404*−/− we crossed several zebrafish of known genotype (*gad1a*−/− or *gad1a* x wild-type) and performed the experiment on three rounds of larvae from these crosses on one day. Embryos were grown in standard egg water (Westerfield, 1993).

To perform the assay, we divided 7 dpf larvae of the desired genotype into six petri dishes each with 15-20 fish and labeled each dish corresponding to the dose that would be assayed. Larvae were allowed to acclimate for 30 min. We remove as much egg water as possible and pour 15 mL of pre-measured PTZ, a known GABA_A_ receptor antagonist, diluted in standard egg water at the following concentrations into the appropriately labeled dish: 0 mM (egg water only control), 1 mM, 2.5 mM, 5 mM, 10 mM and 15 mM (positive control). Once the solution is bath applied to the dishes, we began a timer for 10 minutes and monitored all dishes for abnormal behavior as defined by stage II and stage III seizure-like behavior (Baraban et al., 2005). To control for double counting of responding fish, when we saw a fish that exhibited abnormal behavior, we removed that fish and placed it in a separate dish. At the end of 10 minutes, we counted how many fish responded with stage II or stage III behavior and how many fish did not respond at each treatment group.

### Extracellular Electrophysiology

Zebrafish of the desired genotype were grown to 7 dpf and immobilized with 250 μM of α-bungarotoxin in 1X E3 media with 1 mM HEPES (Westerfield, 1993). Once paralyzed, we moved single larvae to the lid of a 35 mm non-tissue-culture-treated petri dish (Corning Inc., Tewksbury, MA) and oriented the fish laterally. Once properly positioned, we added warm, but not hot, 0.4% agarose in 1X E3 media onto the fish and let it cool for ~2 minutes. We added ~3.5 mL of 1X E3 media to the lid and then inserted a sharp glass pipet microelectrode (15-20 MΩ impedance), loaded with 2-3 μL of normal Ringer’s solution (116 mM NaCl, 2.9 mM KCl, 1.8 mM CaCl_2_, 5.0 mM HEPES, pH 7.2) into the optic tectum (TeO). The optic tectum was chosen to facilitate comparison with previously published data obtained from larval zebrafish (Baraban et al., 2005). A chloride-coated silver wire (0.010” A-M Systems, Inc., Sequim, WA) reference electrode was placed touching the surrounding solution. Field recordings were collected using Molecular Devices’ Axoclamp software and data were digitized at 10 kHz, low-pass filtered at 1 kHz, and analyzed with CLAMPEX 10.4 software (Axon Instruments, Sunnyvale, CA). We performed field recordings from each fish for 20 minutes.

### Calcium imaging with light sheet microscopy

Calcium imaging was performed on a custom-built light sheet microscope (Supplemental Fig. S2). The system is a modified version of the OpenSPIM setup (Pitrone et al., 2013), as described in our previous work (Liu et al., 2019a). The microscope is controlled through a custom-written LabVIEW program using a Dell Precision 5810 Tower with 32GB RAM and a quad-core Intel(R) Xeon(R) E5-1603 v3 processor. We followed the protocol described by Huisken lab (Kaufmann et al., 2012; Weber et al., 2014). Transgenic zebrafish larvae (*elavl3*:GCaMP5g; gad1b:RFP; mitfaw2/w2) at 5 to 7 day post-fertilization (dpf) of development were immobilized using 100μM of alpha-bungarotoxin. The fish were then immersed in a 0.2% agarose solution, and inserted into a 1cm cut FEP tube. The tube was then sealed with 3% agarose gel and sealed with parafilm. Each fish was imaged at approximately the same horizontal plane referenced from the dorsal surface of the tectum (Supplemental Fig. S3) continuously for 2 to 10 minutes under the same laser power (10mW, 99.21 W/cm^2^ at the sample). Imaging data was collected at 33-50 frames per second (fps) for each single channel.

For experiments imaging PTZ-induced neural activity, larvae were treated with 15 mM of PTZ for 40 min before mounting for light-sheet imaging.

### Image Analysis

For frequency analysis, the image time series was first registered using the method described in ref (Guizar-Sicairos et al., 2008). Whole frame average intensities were computed for each image time series. The first 100 frames (4.4 seconds) were omitted from this computation because the LSM excitation light initially activates neural activity in the larvae. Then fluorescence intensity changes (ΔF/F) were calculated using the sliding window method described (Patel et al., 2015; Liu and Baraban, 2019). This method involves finding the median value in the window interval before each data point (*F*_t0_), subtracting the mean (*F_μ_*) of the data points below the median from the original data point(*F*_t0-Δ*t*_), then normalizing by dividing this result by the same mean value. This process is described by the following equation:

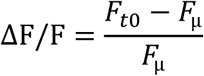

Next, we performed a Fourier Transform on the one-dimensional fluorescence traces to get the frequency spectra. Then we calculated the absolute value of the real part of the frequency spectrum and normalized it to the maximum value of the frequency spectrum. This analysis was performed for three fish each from three different groups (control, PTZ-treated*, gad1b* null).

In order to compare across the changes in neural activity within the optic tectum of the zebrafish larva, we register our image times series to the zebrafish brain atlas (Kunst et al., 2019b) and find the position of the imaged plane in 3D using cross-correlation. Then the masks of regions of interests are downloaded from the brain atlas website (https://fishatlas.neuro.mpg.de/). We apply the non-rigid image registration method (Garyfallidis et al., 2014) to segment these ROIs in our data and extract the signal of each ROI by taking the average of the image intensity in each ROI’s region. Afterward, we calculate the correlation values between these ROIs using the extracted signals.

We adapted the sliding-window framework (Zalesky et al., 2014; Hindriks et al., 2016) to analyze the dynamic connectivity of the regions of interest. Specifically, we use a tapered window of length 20 s. We slide the window in time at a temporal resolution of 3s over the 5-min interval and get a continuous series of snapshots of the ROIs signals. For each snapshot, we calculate a correlation matrix for the ROIs using Pearson correlation. Finally, we get a series of correlation matrices (regions × regions × windows).

We consider two regions to be connected if the corresponding correlation is larger than 0.6. The frequency of connections between the two sides of brains and within each side of the brain are summarized.

## EXPERIMENTAL DESIGN AND STATISTICAL ANALYSES

For the HPLC-ECD experiments with 7 dpf larval samples, we assayed 5 replicates, in this case 5 single larval heads. For the adult samples, we used 5 male and 5 female brains to account for potential gender differences across the genotypes tested. The selection of 5 biological replicates came from discussions with HPLC experts to capture sufficient amounts of data for proper statistics. Results of these experiments were analyzed and plotted using GraphPad Prism (La Jolla, CA). We plotted normalized values from all replicates for each genotype with the mean and standard deviation. We performed a one-way ANOVA to test statistical significance across the groups.

For the PTZ dose response assay, we used 15-20 7 dpf zebrafish per treatment group for each genotype tested. When we assayed a *gad1b* heterozygous incross, we randomly sorted zebrafish into pools of 20 with the assumption that ~5 fish per genotype would be in each treatment group. When we assayed crosses of known genotypes, we used 20 zebrafish per treatment group when assaying the *gad1b* allele and 15 zebrafish per treatment group when assaying the gad1a allele. The *gad1b* heterozygous incross resulted in unequal numbers of each genotype for that experiment, thus the resulting N value is not the same across each genotype. However, each of the 10-minute assays were performed 3 times for each genotype. Taking all experiments into account, we assayed at least 42 larvae for each genotype at each PTZ treatment which provides more than sufficient biological and technical replicates for statistics. Gender is not determined in 7 dpf larvae, so we did not consider sex differences in this experiment. Results of these experiments were plotted using GraphPad Prism (La Jolla, CA). Due to unequal numbers across the genotypes, we plotted the percentages of responding fish of each genotype at each dose with standard deviation across the three trials. We performed no additional statistical analyses on these data.

For the electrophysiology experiments, we used at least ten fish per genotype or treatment group. This is the standard number of replicates used in the field to capture any natural variation that occurs across a population. The raw trace data is reported, so there were no statistical analyses performed on these data.

For the analysis of brain connectivity, eight wild-type fish, nine PTZ treated wildtype fish, seven *gad1b* mutant fish and five PTZ treated *gad1b* mutant fish were analyzed. Using the sliding window framework, we calculated the frequency of the functional connectivity within and between the two half sides of brains for each single run of each fish. To test whether there is any difference of the connectivity frequency between different groups, we applied the Wilcoxon rank sum test (Mann and Whitney, 1947) for each two different groups and calculate the P-values based on the corresponding Mann-Whitney U statistics.

## Results

### Zebrafish homozygous null for *gad1a* or *gad1b* exhibit reduced levels of GABA in the brain and increased neural activity

CRISPR-Cas9 mediated genome editing was used to generate sets of indels in the *gad1a* and *gad1b* genes in zebrafish. Two indels, allele ga2404 for *gad1a* and allele ga2303 for *gad1b,* were predicted null mutations and selected for this study (Figure 1A). Zebrafish heterozygous for mutations in *gad1a* or *gad1b* were crossed to generate homozygous mutant lines. Zebrafish homozygous for *gad1a^ga2404^* or *gad1b^ga2303^* have normal gross body morphology, are viable and fertile.

**Figure 1.**
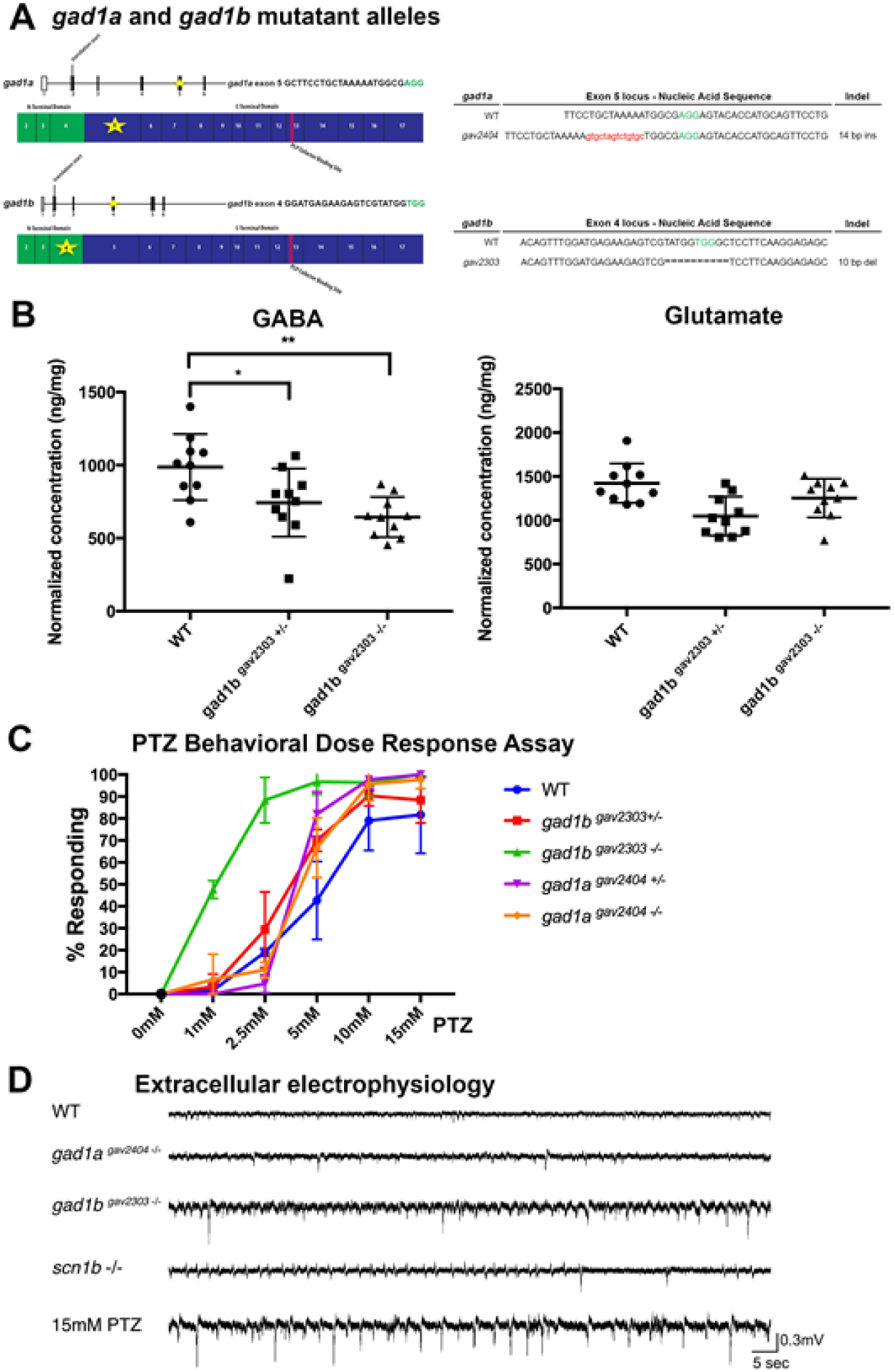
*gad1b* mutant larvae exhibit increased neural activity. (A) Location of CRISPR-Cas generated indels the *gad1a* and *gad1b* genes. (B) GABA and glutamate levels in the brains of *gad1b*^+/+^, *gad1b*^+/−^, and *gad1b^-^*^/-^ fish as measured by HPLC with electrochemical detection. Samples were normalized and run as described previously (Ross and Filipov 2006, Coban and Filipov 2007); N=10 animals/genotype. (C) PTZ dose response assay on 7 dpf *gad1a* and *gad1b* mutant larvae. Fish were sorted into groups of 10-20 per genotype for each treatment group and assayed for stage II and stage III behavioral phenotypes for 10 minutes. Assays were performed three times each. N>42 for each genotype at each dose. (D) Representative traces of extracellular recordings made from 7 dpf *gad1b*^−/−^ mutant larvae compared to wild-type (WT), *gad1a*^−/−^, *scn1b*^−/−^, and WT larvae exposed to 15 mM PTZ. N>10 for each genotype/treatment.

High performance liquid chromatography with electrochemical detection (HPLC-ECD) was used to assess the levels of GABA, glutamine, glutamate, and monoamine neurotransmitters in the brains *gad1a^ga2404^* and *gad1b*^ga2303^ mutant zebrafish compared to wild type fish (Fig. 1B, Supplemental Tables 2-4). Levels of GABA, glutamine and glutamate were assessed separately from serotonin (5-HT), 5-Hydroxyindoleacetic acid (5-HIAA), dopamine (DA), norepinephrine (NE), and 3-Methoxy-4-hydroxyphenylglycol (MHPG) because of different requirements in sample preparation. Amino acid and monoamine neurotransmitter data for adult brains was obtained using 20 individual brains for wild-type, *gad1a*^−/−^ and *gad1b*^−/−^, and 10 brains for *gad1b*^+/−^, with equal numbers of brains for each sex. Brain samples were age and sex matched between genotypes. Brain weights ranged from 1.2 mg to 7.2 mg (mean = 4.2 mg sd = 1.3) for wild-type fish, 0.7 to 4.0 mg (mean = 2.0 mg sd=0.8) for *gad1a*^−/−^, 1.1 mg to 7.4 mg (mean = 3.9 mg sd = 2.0) for *gad1b*^+/−^, and 1.0 to 7.0 mg (mean = 3.5 mg sd=1.5) for *gad1b^−/−^.* Decreased levels of GABA were measured in the adult brains of fish mutant for *gad1a* or *gad1b.* The average normalized concentration of GABA in adult wild-type brain was determined to be 921 ng/mg of tissue (sd = 261) compared to 678 ng/mg of tissue (sd = 256) for fish homozygous mutant for *gad1a^ga2304^,* 743 ng/mg of tissue (sd=221) for fish heterozygous mutant for *gad1b*^ga2303^, and 652 ng/mg of tissue (sd = 135) for fish homozygous mutant for *gad1b^ga2303^* (Fig. 1B). Zebrafish heterozygous for *gad1a*^ga2404^ were not tested. Male and female fish exhibited comparable levels of GABA within genotypes (data not shown). Comparable levels of glutamate, glutamine, serotonin (including 5-Hydroxyindoleacetic acid), dopamine, and norepinephrine (including 3-Methoxy-4-hydroxyphenylglycol) were measured in the adult brains of all genotypes (Fig. S4). These data support that *gad1a^ga2404^* and *gad1b*^ga2303^ are functional null mutations for *gad1a* and *gad1b*, respectively.

Determining neurotransmitter levels in 7 dpf larvae was more challenging because of the small amounts of tissue obtained from each animal. It was not practicable to dissect out the brains; therefore, for these experiments, heads were dissected just posterior to the otocyst and anterior to the pectoral fins as these were clearly identifiable morphological features that could be easily used to guide cuts. Cuts were made at an angle to avoid the swim bladder and yolk and to try to minimize the amounts of non-neural tissues included in the sample. The average amount of tissue collected from single 7 dpf larval zebrafish heads was 0.20 mg (sd = 0.14), with no significant differences between genotypes (data not shown). HPLC-ECD analyses were performed using pools of 10 larval heads for each genotype (N = 5 pooled samples / dataset). The average amount of tissue in each pool was 2.52 mg (sd = 0.46). Decreased GABA levels were measured for *gad1b*^−/−^ larvae but not *gad1a*^−/−^ larvae (Figure S5). The average normalized concentration of GABA in wild-type larvae was determined to be 20.9 nM/mg of tissue (sd = 2.3) compared to 8.2 nM/mg of tissue for *gad1b*^−/−^ larvae (sd = 1.7). The average normalized concentration of GABA in *gad1a*^−/−^ larvae was determined to be 24.4 nM/mg of tissue (sd = 4.2). Comparable concentrations of glutamate and glutamine were measured across all genotypes (Supplemental Table 4, Figure S5). Monoamine neurotransmitters were not assessed for 7 dpf larvae. These results indicate that Gad1b enzymatic activity produces a large percentage of the GABA found in the 7 dpf larval zebrafish brain.

Larval zebrafish harboring mutations in *gad1a* or *gad1b* are sensitive to pharmacological perturbations in GABA signaling (Figure 1C). Larval zebrafish of different *gad* genotypes at 7 days of development (7 dpf) were exposed for 10 min to different doses of the proconvulsive compound pentylenetetrazole (PTZ), which is a non-competitive GABA antagonist, and scored for the first appearance of Stage II or Stage III motor behaviors during the 10-minute exposure interval. As first described by Baraban (2005), Stage II behavior is characterized by a rapid, tight circular swim trajectory and Stage III behavior is characterized by a loss of posture and mobility for 1-3 s. During these experiments, all larvae exhibited normal swimming behavior in the absence of PTZ and in a visually and sonically neutral environment. Larvae mutant for *gad1a* or *gad1b* were more sensitive to PTZ than were wild-type larvae (Figure 1B). Notably, almost half of the *gad1b^−/−^* larvae and about 7% of the *gad1a*^−/−^ larvae, but none of the wild-type larvae, exhibited Stage II/III motor behavior when exposed to 1 mM PTZ. These results suggest that GABA levels are reduced in the *gad1a^ga2404^* and *gad1b*^ga2303^ mutant lines, with the greater reduction associated with *gad1b*^ga2303^.

Extracellular electrophysiology was used to measure local field potentials in the optic tecta of *gad1a^ga2404^* or *gad1b*^ga2303^ homozygous mutant 7 dpf larval zebrafish to compare to those recorded from wild-type larvae, wild-type larvae exposed to 15 mM PTZ, and larval zebrafish homozygous mutant for *scn1ab* (Figure 1D). Ten larvae were used for each condition. Larvae homozygous mutant for *gad1a^ga2404^* or *gad1b*^ga2303^ exhibited increased electrographic activity in the optic tectum compared to wild-type larvae, with the larger increase associated with mutations in *gad1b*. The pattern of activity observed in *gad1b*^−/−^ larvae was most similar to that observed for wild-type larvae treated with 15 mM PTZ than for 7 dpf larvae homozygous mutant for *scn1a*. The *scn1a* gene encodes for a subunit of the Nav1.1 voltage-gated sodium channel (Grone et al., 2017). Notably, both *gad1b* mutant larvae and wild-type larvae treated with 15 mM PTZ exhibited increased amplitude of high-frequency discharges punctuated with high-amplitude, low frequency discharges. These results show that *gad1b* gene function plays a more significant role than *gad1a* for normal neural signaling in the optic tectum. Additionally, these results indicate that GABAergic neurons in the optic tectum mediate activity in a relatively large set of neural circuits with similar properties, the existence of which is revealed in the similar electrographic features observed for both *gad1b* mutant larvae and wild-type larvae treated with 15 mM PTZ, but not in *scn1a* mutant larvae.

### *gad1b and gad2* gene are expressed by most GABAergic neurons in the developing zebrafish optic tectum

Expression of *gad1a*, *gad1b* and *gad2* in the optic tectum was assessed by mRNA *in situ* hybridization at 3 dpf and 5 dpf both in whole-mount and in horizontal sections cut through the optic tectum (Fig. 2). Connections between retina and optic tectum become functional between 3-4 dpf (Stuermer, 1988; Burrill and Easter, 1994; Easter and Nicola, 1996, 1997; Niell and Smith, 2005). By 5 dpf, larvae track and capture prey indicating a functional visual system (Niell and Smith, 2005). Anatomically, the larval tectum has two distinct regions, one composed of neuronal cell bodies, known as the stratum periventriculare (SPV), and the other a superficial neuropil that is organized into layers (Fig. 2 H,J). The neuropil contains the dendrites and axons of tectal neurons, a sparse mixture of tectal interneurons and afferent axons arriving at the tectum, mostly from the retina (Nevin et al., 2010; Kunst et al., 2019a).

**Figure 2.**
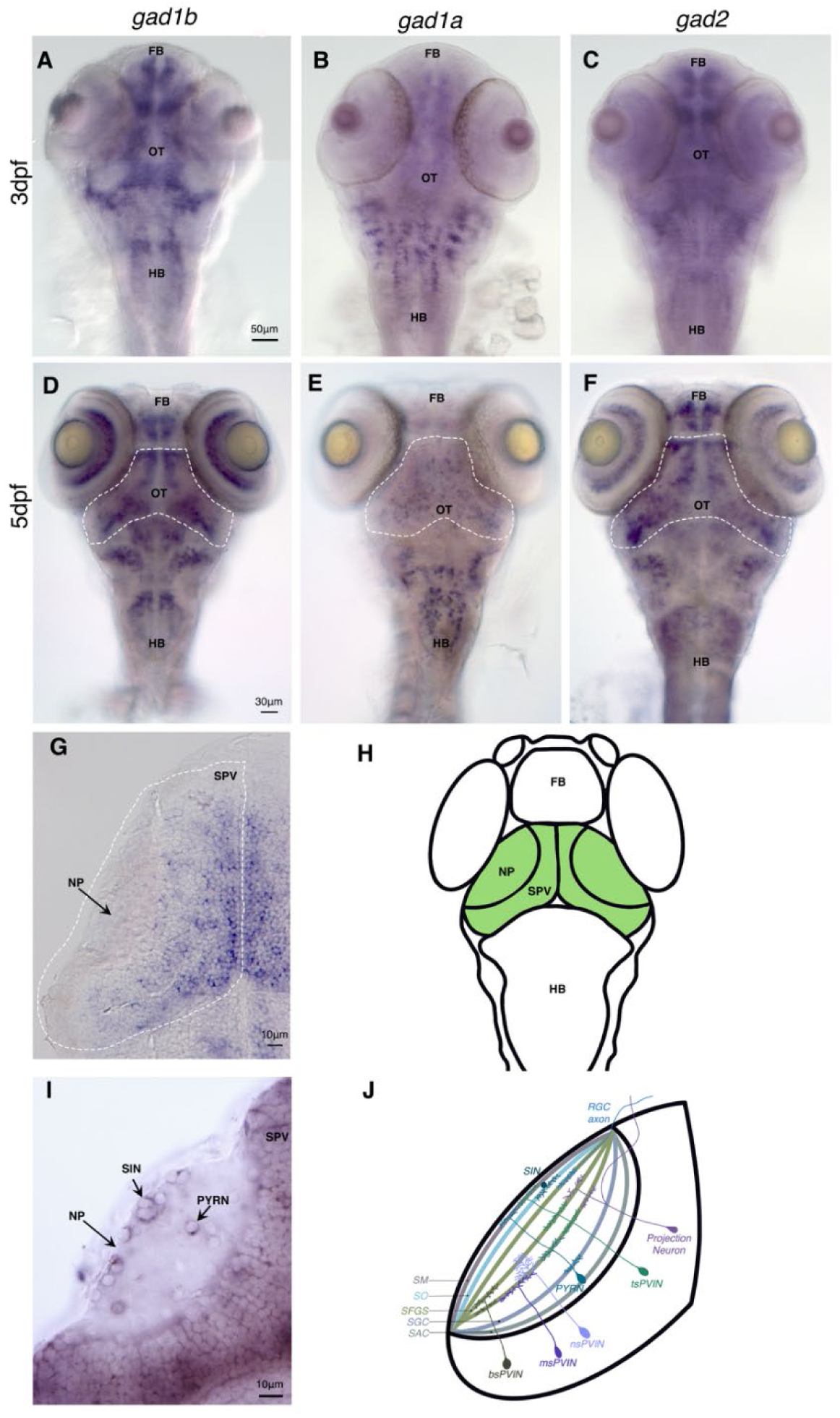
*gad1a* and *gad1b* expression in the larval zebrafish brain. (a-c). 3dpf RNA whole mount *in situ* hybridization of *gad1b*, *gad1a*, and *gad2*. (d-f) 5dpf whole mount *in situ* hybridization. Dashed line outlining optic tectum. (g). Section of *gad1b* expression in the optic tectum. Dashed line outlining hemitectum. (h). Schematic of larval zebrafish highlighting optic tectum in green. (i) Section of 5dpf *in situ* hybridization in the optic tectum. (j) schematic of a hemitectum showing examples of cell bodies and their neurite patterns in the neuropil. FB: forebrain, OT: optic tectum, HB: hindbrain, NP:Neuropil, SPV: stratum periventricular

At 3 dpf, *gad1a*, *gad1b* and *gad2* exhibit distinct patterns of expression in the developing brain (Figure 2). *gad1a* is predominantly expressed by cells in a longitudinal domain adjacent to the ventral midline that extends from the hypothalamus through the tegmentum and also in clusters of cells in the hindbrain. No appreciable expression is detected in neurons in the telencephalon or in the optic tectum. *gad1b* is predominantly expressed by clusters of cells in the telencephalon, diencephalon, optic tectum, and hindbrain. *gad2* expression overlaps with *gad1a* and *gad1b* in the brain, but *gad1a* and *gad1b* are mostly expressed by separate cells in the forebrain, midbrain and hindbrain.

In the larval optic tectum at 5 dpf, *gad1b* and *gad2* are expressed by cells in the neuropil and SPV. Notably, *gad1b* and *gad2* are expressed by neurons with cell bodies located superficially in the neuropil between the stratum opticum (SO) and the stratum fibrosum et griseum superficiale (SFGS) laminae, likely superficial interneurons (SINs), and also in a subset of neurons with cell bodies located in the deep layers of the neuropil. These latter neurons are likely GABAergic pyramidal neurons (PyrNs) (Nevin et al., 2010; Kunst et al., 2019a; DeMarco et al., 2020). *gad1b* and *gad2* expression in the SPV is detected in clusters of cells, some of which are likely periventricular interneurons (PVINs). The SPV is comprised of radial glia and at least 19 different types of neurons (Nevin et al., 2010; Robles et al., 2011; Kunst et al., 2019a; DeMarco et al., 2020). Of these, GABAergic PVINs make up approximately 20% of the neurons in the SPV (Scott et al., 2007; Scott and Baier, 2009). *gad1a* transcripts were detected mostly in cells in the SPV.

As a second means of assessing *gad1b* expression in the tectum, reporter gene expression was assessed in larvae stably transgenic for the *gad1b*-reporter transgene TgBAC[*gad1b*:*loxP-DsRed-loxP-GFP*] (Satou et al., 2013). This reporter drives DsRed expression in putative SINs and PyrNs in the tectal neuropil and in neurons in the SPV (Figure S6).

Together, these data indicate that null mutations in *gad1b* should result in a reduction in GABA in neurons involved in processing visual information in the optic tectum.

### *gad1b* null larvae exhibit increased neural activity in the optic tectum

To assess spatiotemporal patterns of neural activity in the optic tectum, calcium imaging was performed using 5 dpf larval zebrafish stably harboring Tg(*elavl3:GCaMP5g*), which drives expression in most if not all neurons in the optic tectum (Ahrens et al., 2013a; Ahrens et al., 2013b). Imaging was performed using a custom built light sheet microscope (LSM), the details of which have been published elsewhere (Liu et al., 2019b). Larvae at 5 dpf were chosen because larvae at this stage of development can track and capture prey and avoid predators (Niell and Smith, 2005), which are behavioral indicators of a functional visual system, and, in our hands, are more easily imaged than larvae at older stages of development even with pigment suppression. It should be noted that the size of the visual receptive field has been reported to increase between the stages of 4 dpf to 6 dpf and then reduce by 8-9 dpf (Zhang et al., 2011). Thus, imaging at 5 dpf captures the behavior of neuronal assemblies during the period in which the tectal circuitry is undergoing developmentally and functionally driven refinement. A minimum of five larvae were used for each condition with two to three recordings made from each larvae (*gad1b*^+/+^, *gad1b*^+/+^ exposed to 15 mM PTZ, *gad1b*^−/−^, and *gad1b*^−/−^ exposed to 15 mM PTZ).

Under the conditions used for these experiments, *gad1b*^+/+^ larvae exhibited intermittent increases in GCaMP5g fluorescence predominantly in the neuropil of the anterior tectum and within the superficial neuropil layers (Figure 3A). A typical example is shown by plotting the relative standard deviation (RSD) of each pixel as measured over 2 min at 46 fps. Calcium activity typically increased in the anterior tectum and propagated anterior to posterior within the SO or SFGS (data not shown). Activity was largely independent for each hemi-tecta. An increase in GCaMP5g fluorescence was detected in cell bodies in the SPV, but the largest relative changes in GCaMP5g fluorescence occurred in the neuropil, which is consistent with previous observations that information processing in the teleost tectum appears to take place predominantly, if not exclusively, in the neuropil (Kinoshita et al., 2002; Nevin et al., 2010). Spectral analysis revealed that fluctuations in the GCaMP5g signal occurred predominantly at 3.84 Hz, 7.63 Hz and 8.20 (Figure 3D).

**Figure 3.**
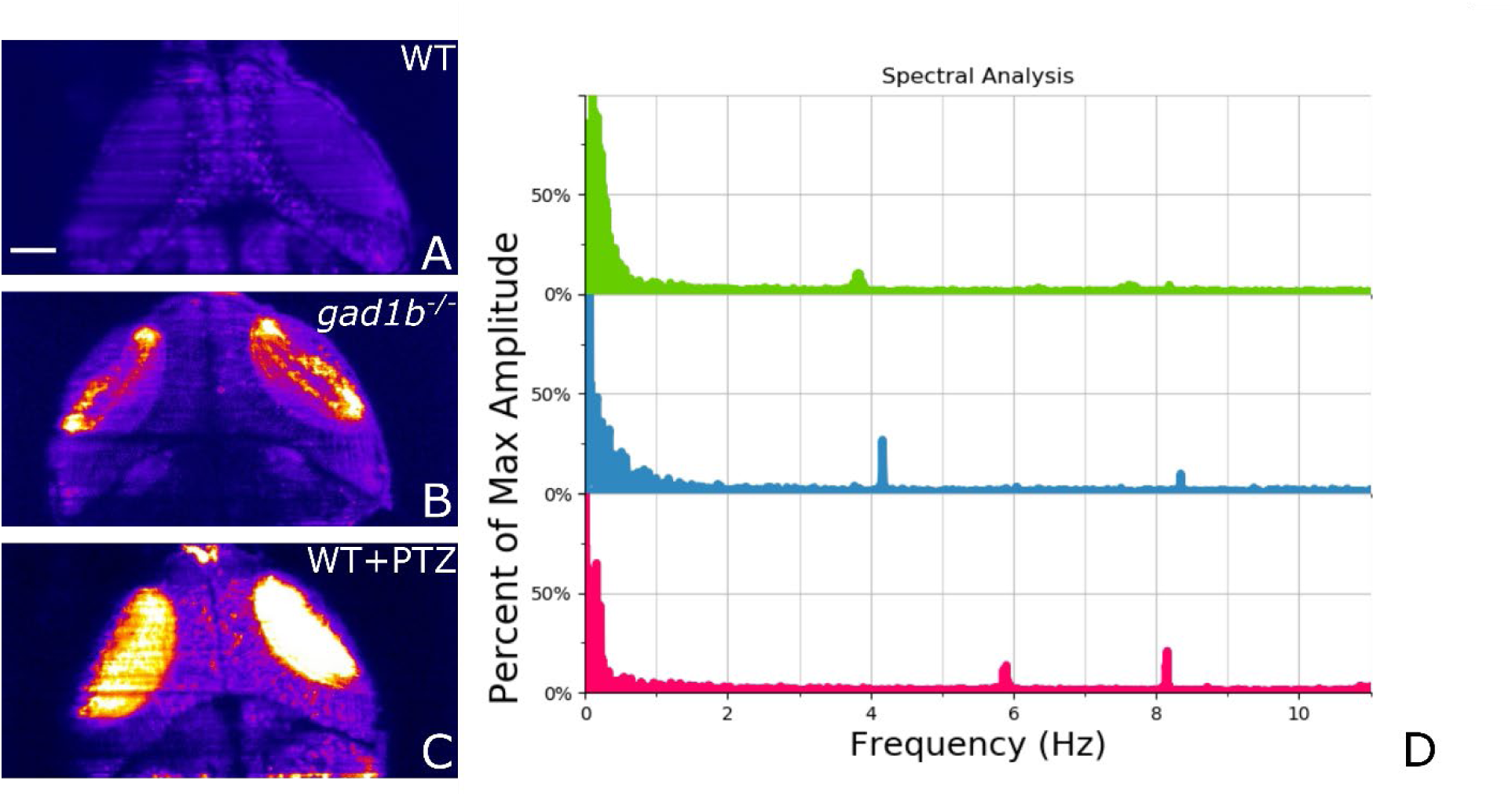
(A)-(C) Plot of the standard deviation over the mean of the GCaMP signal for Wild-type, gad1b−/− mutant, and wild-type treated with PTZ. Analysis is done using the first 100 frames with imaging speed of 25fps (~4 seconds) of each recording. (D) Frequency analysis of 10-minute recordings with photobleaching corrected in deltaF calculations. Wild-type, *gad1b*^−/−^ and, PTZ treated fish each show distinct frequencies in the temporal response. The relevant peaks are 3.84Hz, 7.63Hz, and 8.20Hz for wildtype. The scale bar in (A) is 50 microns.

*gad1b* mutant larvae exhibited changes in both spatial and temporal aspects of the Ca^2+^ signals relative to *gad1b* normal larvae (Figure 3B). Like *gad1b*+/+ larvae, changes in GCaMP5g fluorescence typically initiated in the anterior tectum and propagated posteriorly within neuropil layers, however, there was an expansion in the amount of superficial neuropil that exhibited increased GCaMP5g fluorescence, especially in the areas surrounding putative SINs. In these embryos, increased GCaMP5g fluorescence was often detected in cell bodies in the SPV, typically in concert with increased fluorescence in discrete areas of the neuropil. Spectral analysis revealed fluctuations in the GCaMP5g signal were occurring predominantly at 4.16 and 8.35 Hz (Figure 3D).

To assess the impact of an acute reduction in GABAergic signaling, *gad1b*^+/+^ larvae were bath exposed to 15 mM PTZ. These larvae exhibited widespread changes in GCaMP5g fluorescence that included all regions of the neuropil and often cell bodies in the SPV (Fig. 3C). In these larvae, activity often propagated posterior to anterior in the tectum (data not shown). Spectral analysis revealed fluctuations in the GCaMP5g signal were occurring predominantly at 5.91 and 8.15 Hz (Fig. 3D). The PTZ data indicates that GABA is acting predominantly as an inhibitory neurotransmitter in the tecta of 5 dpf larvae and that GABAergic circuits are governing information flow through tectal circuits.

Regional activity within the optic tectum was assessed for individual larvae (Figures 4, 5) and then compared within experimental groups and across conditions (Figure 6). In wild-type larvae, in the absence of PTZ, activity was largely restricted to the neuropil (Figs. 4A, 5A). Within the neuropil of a hemitecta, correlated spikes of calcium activity were observed in the regions of the SO and SFGS, but not all activity in SFGS correlated with that of the SO. The timing of activity in the neuropil adjacent to the SPV usually correlated with that of the SFGS. Activity of the left and right hemitecta appeared to be mostly independent of each other. Exposure of wild-type larvae to PTZ resulted in concomitant spikes of calcium activity across all layers of the neuropil as well as within the SPV (Fig. 4C,H; 5B,E,H). Interestingly, exposure of wild-type fish to PTZ resulted no correlation between the left and right hemitecta (Fig. 4H). Like wild-type, larvae null for *gad1b* exhibited correlated activity between the SO and SFGS, but not all activity in the SFGS correlated with the SO, and changes in activity in the SFGS often correlated with changes in the neuropil adjacent to the SPV. Unlike wild-type larvae, concomitant spikes of calcium activity were observed in both the SFGS and in neuronal soma in the SPV (Fig. 4C,G; 5C,F,I). Exposure of gad1b−/− larvae to PTZ resulted in increased, correlated activity in both the left and right hemitecta (Fig. 4i).

**Figure 4.**
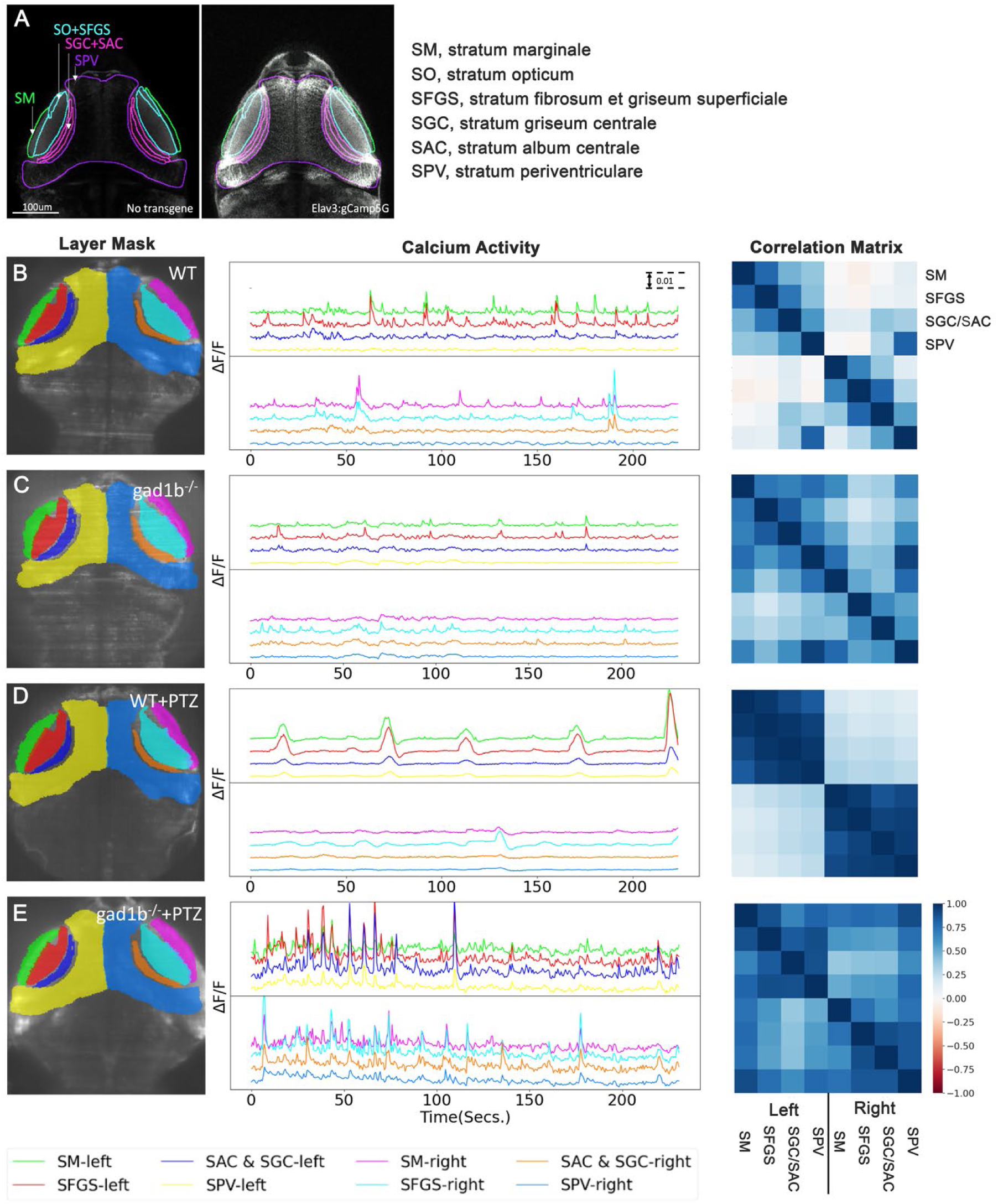
Calcium activity by tectal region. (A) Regions of the optic tectum. (B) Neural activity in the wild-type optic tectum. (C) Neural activity in the *gad1b*^−/−^ tectum. (D) Neural activity in wild-type larva treated with PTZ. (E) Neural activity in a *gad1b*^−/−^ larva treated with PTZ. SM, stratum marginale; SO, stratum opticum; SFGS, stratum fibrosum et griseum superficiale; SGC, stratum griseum centrale, SAC, stratum album centrale

**Figure 5.**
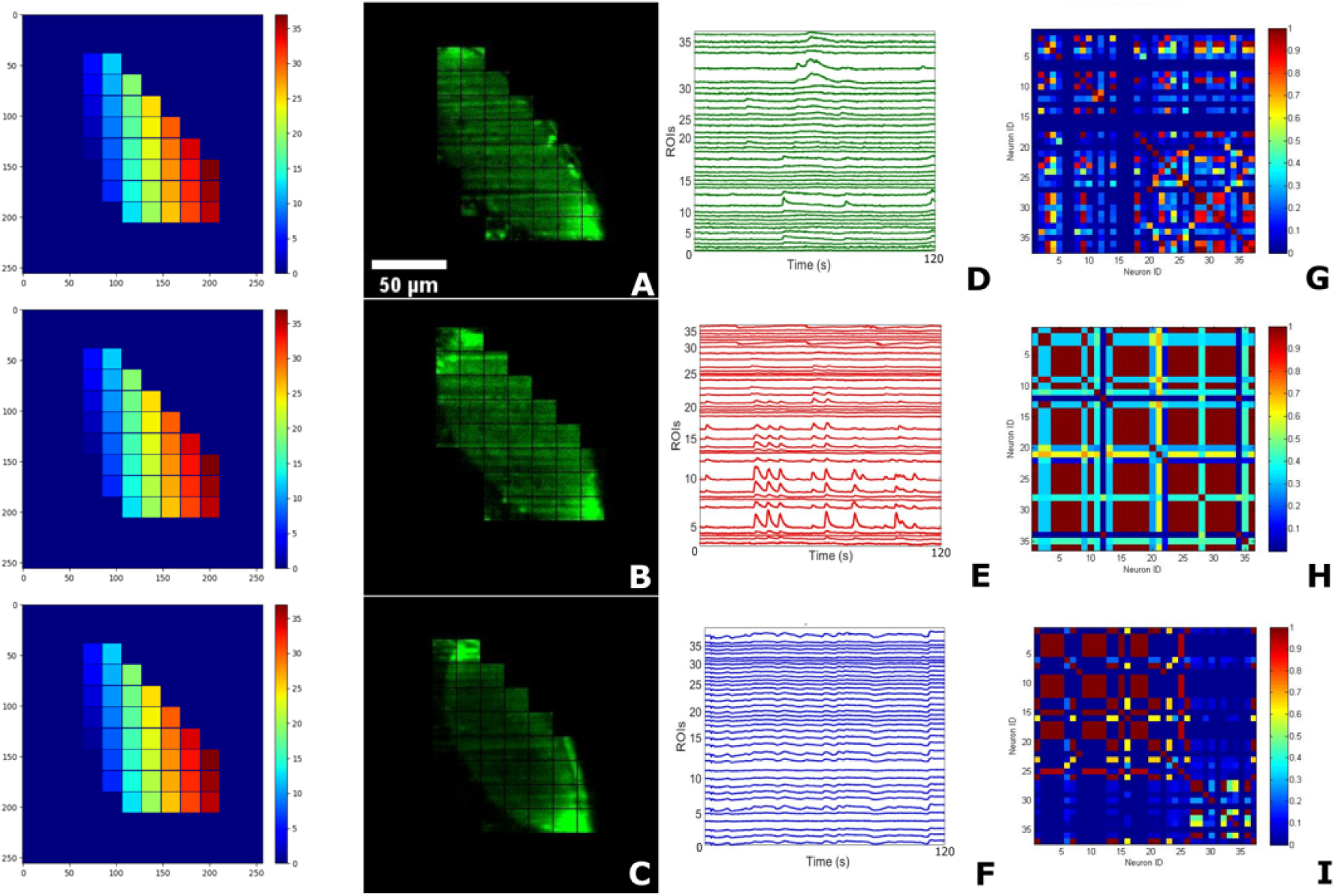
Optic Tectum Analysis. nalysis is similar to in Fig. 4 but now measuring activity in different regions of the optic tectum. (A)-(C) Images of the right tectum in wildtype, PTZ treated and *gad1b*^−/−^ zebrafish larva. (D)-(F) activity in the different regions. (G)-(I) Corresponding correlation matrices.

**Figure 6.**
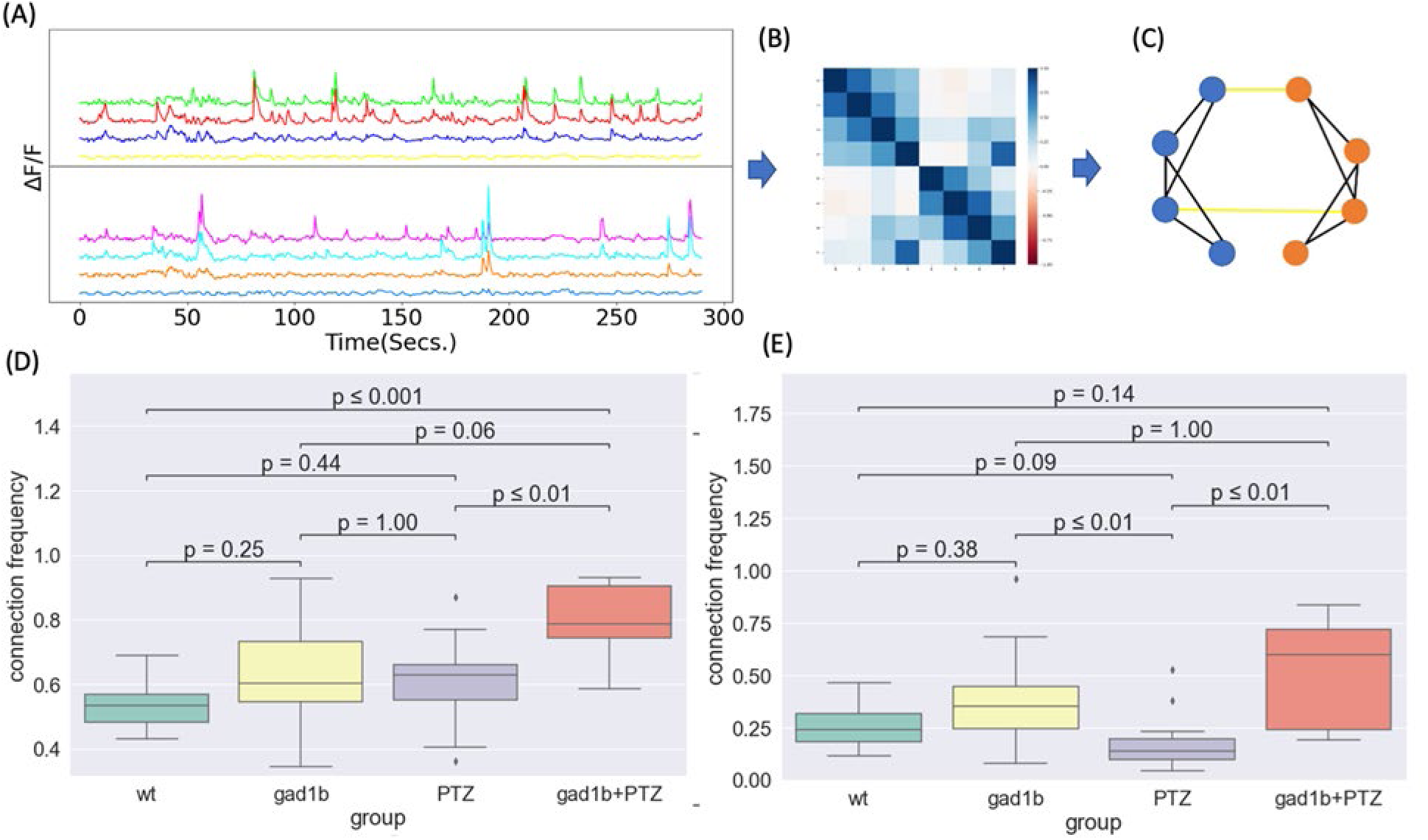
Connectivity analysis. (a) The correlation of activity is calculated using a 20s sliding window with a 3s step size. (b) The correlation matrix between the different regions. (c) A connectivity map is created between the different regions by counting regions as connected for a correlation greater than 0.5. (d) The connectivity between regions on the same side of the brain is compared between the different types. (e) The connectivity between regions on different sides of the brain. N = 5 fish per group with 2-3 recordings per fish.

Assessment of the connection frequency, Fig. 6, revealed that *gad1b* mutant larvae exhibited greater variance than wild-type fish for connectivity within a hemitecta and between the left and right tecta. Interestingly, the connectivity in *gad1b* mutant larvae exposed to PTZ was significantly higher than that of wild-type larvae exposed to PTZ under the same conditions both within a hemitecta and between the left and right sides of the tectum.

## Discussion

Our study investigated the impact of genetically reduced GABAergic signaling on the network dynamics of neuronal populations in the optic tectum of larval zebrafish. We combined genetics and high-speed light sheet imaging of calcium dynamics to investigate the neural response of zebrafish larvae null for the *gad1b* gene compared to wild-type larvae in the presence and absence of PTZ. We used a cross-correlation analysis between different brain regions to indicate potential connectivity. Comparing connectivity data within and between experimental groups revealed that *gad1b* mutant fish exhibited increased connectivity within and between hemitecta compared to wild-type larvae.

Zebrafish have three *gad* genes known as *gad1a*, *gad1b*, and *gad2*. The *gad1a* and *gad1b* genes are paralogs, which likely arose due to a gene duplication event. By mRNA *in situ* hybridization, we showed that cells in the forebrain, inner-nuclear layer of the retina, optic tectum, and hindbrain of larval zebrafish express *gad1b*. At the same developmental time points, strong *gad1a* expression was observed mainly in the hindbrain, with some expression detected in the midbrain and forebrain. These results suggest that specific GABAergic neurons in the zebrafish brain express *gad1a* and *gad1b* and that the two genes are subject to different regulatory inputs. We are still determining if some neurons in the brain co-express *gad1a* and *gad1b*. As predicted by differences in the expression patterns of the two genes, fish mutant for *gad1b* exhibited a more significant decrease in brain GABA levels, a higher sensitivity to PTZ, and a larger increase in baseline neural activity in the optic tectum than did fish mutant for *gad1a*. Cells null for either *gad1a* or *gad1b* likely produce GABA through the action of *gad2* but at lower levels than wild-type cells.

The premise of this study was that genetic reduction of GABA synthesis would result in altered functional connectivity in the developing optic tectum of the zebrafish. In vertebrates, neural circuits are constructed via activity-independent mechanisms and refined by activity-dependent mechanisms. GABAergic signaling has been shown to act as a trophic factor to regulate cell proliferation, neural migration, neurite growth, and synapse formation, as well as mediate neural activity (Ganguly et al., 2001; Kriegstein and Owens, 2001; Ben-Ari, 2002; Manent et al., 2005; Represa and Ben-Ari, 2005; Akerman and Cline, 2007; Ben-Ari et al., 2007). Therefore, perturbations in GABAergic signaling during development can generate different neurological phenotypes depending on when, where, and how signaling is altered. Reduced GABAergic signaling during development has been reported to alter neuron numbers in the brain. In mice, focal pharmacological inhibition of GABAergic signaling in the somatosensory cortex resulted in decreased cell death and increased numbers of neurons in the somatosensory cortex (Duan et al., 2020). A similar study in zebrafish showed that an increase in activity in the brain leads to a decrease in total neuron numbers but an increase in the excitatory-to-inhibitory cell ratio (Brenet et al., 2019). Although the impact of reduced GABAergic signaling on cell survival differs between the two studies, the net effect for both was to shift the excitatory/inhibitory balance in the young brain. An increase in activity would lead to various forms of synaptic plasticity, including long-term potentiation of synaptic responses and subsequent stable alterations in neural networks (Holmes and Ben-Ari, 2001).

Our experiments showed that loss of function mutations in *gad1b* resulted in increased coordinated activity in the larval optic tectum that was different from that observed by acute perturbation of GABAergic signaling by exposure to PTZ. The activity observed in *gad1b* null larvae likely reflects changes in the architecture and information processing of tectal microcircuits. Architecturally, our data suggest that *gad1b* mutant larvae retained more synaptic connections than those present in wild-type larvae. The most substantial support for this idea comes from the different activity phenotypes associated with PTZ exposure. Whereas exposure to wild-type larvae resulted in synchronized activity within either the left or right hemitecta, exposure to *gad1b* null larvae resulted in synchronized activity in both the left and right tecta.

Given that PTZ acts the same way in both wild-type and *gad1b* mutant genotypes, the differences in activity elicited by PTZ exposure likely reflect differences in the geometry of the neuronal networks in wild-type compared to *gad1b* mutant larvae. The spatiotemporal pattern of PTZ-induced activity observed in *gad1b* null larvae is consistent with increased connections in the *gad1b* null tectum relative to the wild-type tectum. An increase in connectivity, measured by magnetic resonance imaging (MRI), has been reported for some children with seizure disorders (Radmanesh et al., 2020; Banerjee et al., 2021).

Information processing is also likely altered in *gad1b* null larvae. The tectum processes spatial information in the visual field, including object location and movement (Gahtan et al., 2005; Del Bene et al., 2010; Nevin et al., 2010). RGC axons enter the tectal neuropil from the anterior side and arborize at one of six retinorecipient laminae (Xiao et al., 2005; Xiao and Baier, 2007; Robles et al., 2013; Robles et al., 2014; Kunst et al., 2019a). In fish that have developed normally, information processing mainly occurs in the tectal neuropil (Kinoshita et al., 2002; Kinoshita and Ito, 2006), with spatial filtering achieved by feedforward inhibition (Del Bene et al., 2010). A reduction in GABAergic signaling, especially in SIN neurons, is expected to result in increased activity in the neuropil in response to a visual stimulus, with deeper layers of the neuropil exhibiting greater activity (Del Bene et al., 2010). Consistent with this expectation, *gab1b* mutant larvae exhibited more widespread activity than in wild-type larvae but less than in wild-type larvae exposed to PTZ.

Our results indicate that a reduction in GABAergic signaling during early brain development results in increased connectivity concomitant with reduced precision of information flow within the brain. In the case of larval zebrafish, our results predict that *gad1b* mutant larvae will exhibit reduced visual acuity.

## Code availability

The code used in this study is available at: https://github.com/Knerlab/neural-activity-analysis

## Data availability

The data used in this study is available at: https://www.ebi.ac.uk/biostudies/bioimages/studies/S-BIAD1200

Because a 10-minute recording at 46 frames per second requires 54 GB of storage, the whole dataset – 73 recordings of 2 to 10 minutes – requires over 2TB of storage. We have chosen to upload 100 frames from each recording. Complete recordings and datasets are available upon reasonable request.

## Acknowledgements

The authors wish to thank Dr. Scott Baraban for generously providing us with the line of *scn1b* mutant zebrafish. The authors thank Brian Condie, Andrew Sornborger, Lindsey Beebe, Madison Grant, Karl Kudyba, and Ashley Rasys for helpful discussion on this project. The authors also wish to acknowledge the UGA Honors Program and the Center for Undergraduate Research Opportunities which supported Mr. J. Branson Byers and Ms. Hannah Schriever in the form of CURO Summer Fellowships and CURO Research Assistantships. This work was supported by the National Institutes of Health (Grant No. R01NS090645 to JDL, PAK and F31NS115496 to YL) and the National Science Foundation (Grant No. 1350654 to PAK). YC and PM were partially supported by National Science Foundation grants DMS-1925066, DMS-1903226, DMS-2124493, DMS-2311297, DMS-2319279, DMS-2318809, and National Institutes of Health grant R01GM152814.

## References

Abrahams BS, Geschwind DH (2008) Advances in autism genetics: on the threshold of a new neurobiology. Nature reviews genetics 9:341–355.

Ahrens MB, Huang KH, Narayan S, Mensh BD, Engert F (2013a) Two-photon calcium imaging during fictive navigation in virtual environments. Front Neural Circuits 7:104.

Ahrens MB, Orger MB, Robson DN, Li JM, Keller PJ (2013b) Whole-brain functional imaging at cellular resolution using light-sheet microscopy. Nat Methods 10:413–420.

Akerman CJ, Cline HT (2007) Refining the roles of GABAergic signaling during neural circuit formation. Trends Neurosci 30:382–389.

Antinucci P, Hindges R (2016) A crystal-clear zebrafish for in vivo imaging. Sci Rep 6:29490.

Arevian AC, Kapoor V, Urban NN (2008) Activity-dependent gating of lateral inhibition in the mouse olfactory bulb. Nat Neurosci 11:80–87.

Asada H, Kawamura Y, Maruyama K, Kume H, Ding RG, Kanbara N, Kuzume H, Sanbo M, Yagi T, Obata K (1997) Cleft palate and decreased brain gamma-aminobutyric acid in mice lacking the 67-kDa isoform of glutamic acid decarboxylase. Proc Natl Acad Sci U S A 94:6496–6499.

Asada H, Kawamura Y, Maruyama K, Kume H, Ding R, Ji FY, Kanbara N, Kuzume H, Sanbo M, Yagi T, Obata K (1996) Mice lacking the 65 kDa isoform of glutamic acid decarboxylase (GAD65) maintain normal levels of GAD67 and GABA in their brains but are susceptible to seizures. Biochemical and biophysical research communications 229:891–895.

Avitan L, Pujic Z, Molter J, Van De Poll M, Sun B, Teng H, Amor R, Scott EK, Goodhill GJ (2017) Spontaneous Activity in the Zebrafish Tectum Reorganizes over Development and Is Influenced by Visual Experience. Curr Biol 27:2407–2419 e2404.

Banerjee S, Dong M, Lee MH, O’Hara N, Juhasz C, Asano E, Jeong JW (2021) Deep Relational Reasoning for the Prediction of Language Impairment and Postoperative Seizure Outcome Using Preoperative DWI Connectome Data of Children With Focal Epilepsy. IEEE Trans Med Imaging 40:793–804.

Baraban SC (2005) Modeling Epilepsy and Seizures in Developing Zebrafish Larvae. In: Models of Seizures and Epilepsy, 1 Edition (Pitkänen A, Schwartzkroin PA, Moshé SL, eds), pp 189–198. 30 Corporate Drive, Suite 400, Burlington, MA 01803: Elsevier Academic Press.

Baraban SC, Dinday MT, Hortopan GA (2013) Drug screening in Scn1a zebrafish mutant identifies clemizole as a potential Dravet syndrome treatment. Nat Commun 4:2410.

Baraban SC, Taylor MR, Castro PA, Baier H (2005) Pentylenetetrazole induced changes in zebrafish behavior, neural activity and c-fos expression. Neuroscience 131:759–768.

Barker AJ, Baier H (2015) Sensorimotor decision making in the zebrafish tectum. Curr Biol 25:2804–2814.

Beck S, Hallett M (2011) Surround inhibition in the motor system. Exp Brain Res 210:165–172.

Ben-Ari Y (2002) Excitatory actions of gaba during development: the nature of the nurture. Nature reviews Neuroscience 3:728–739.

Ben-Ari Y (2006) Seizures Beget Seizures: The Quest for GABA as a Key Player. 18:135-144.

Ben-Ari Y, Gaiarsa JL, Tyzio R, Khazipov R (2007) GABA: a pioneer transmitter that excites immature neurons and generates primitive oscillations. Physiological reviews 87:1215–1284.

Brenet A, Hassan-Abdi R, Somkhit J, Yanicostas C, Soussi-Yanicostas N (2019) Defective Excitatory/Inhibitory Synaptic Balance and Increased Neuron Apoptosis in a Zebrafish Model of Dravet Syndrome. Cells 8.

Bruford EA, Braschi B, Denny P, Jones TEM, Seal RL, Tweedie S (2020) Guidelines for human gene nomenclature. Nat Genet 52:754–758.

Bult CJ, Blake JA, Smith CL, Kadin JA, Richardson JE, Mouse Genome Database G (2019) Mouse Genome Database (MGD) 2019. Nucleic Acids Res 47:D801–D806.

Burgstaller J, Hindinger E, Gesierich B, Baier H (2019) Light-sheet imaging and graph-theoretical analysis of antidepressant action in the larval zebrafish brain network. bioRxiv.

Burrill JD, Easter SS, Jr (1994) Development of the retinofugal projections in the embryonic and larval zebrafish (*Brachydanio rerio*). J Comp Neurol 346:583–600.

Chatron N et al. (2020) Bi-allelic GAD1 variants cause a neonatal onset syndromic developmental and epileptic encephalopathy. Brain 143:1447–1461.

Coban A, Filipov NM (2007) Dopaminergic toxicity associated with oral exposure to the herbicide atrazine in juvenile male C57BL/6 mice. J Neurochem 100:1177–1187.

Condie BG, Bain G, Gottlieb DI, Capecchi MR (1997) Cleft palate in mice with a targeted mutation in the gamma-aminobutyric acid-producing enzyme glutamic acid decarboxylase 67. Proc Natl Acad Sci U S A 94:11451–11455.

de Curtis M, Avoli M (2016) GABAergic networks jump-start focal seizures. Epilepsia 57:679–687.

Del Bene F, Wyart C, Robles E, Tran A, Looger L, Scott EK, Isacoff EY, Baier H (2010) Filtering of visual information in the tectum by an identified neural circuit. Science 330:669–673.

DeMarco E, Xu N, Baier H, Robles E (2020) Neuron types in the zebrafish optic tectum labeled by an id2b transgene. J Comp Neurol 528:1173–1188.

Duan ZRS, Che A, Chu P, Modol L, Bollmann Y, Babij R, Fetcho RN, Otsuka T, Fuccillo MV, Liston C, Pisapia DJ, Cossart R, De Marco Garcia NV (2020) GABAergic Restriction of Network Dynamics Regulates Interneuron Survival in the Developing Cortex. Neuron 105:75–92 e75.

Easter SS, Jr., Nicola GN (1996) The development of vision in the zebrafish (Danio rerio). Dev Biol 180:646–663.

Easter SS, Jr., Nicola GN (1997) The development of eye movements in the zebrafish (Danio rerio). Dev Psychobiol 31:267–276.

Erlander MG, Tobin AJ (1991) The structural and functional heterogeneity of glutamic acid decarboxylase: a review. Neurochem Res 16:215–226.

Erlander MG, Tillakaratne NJK, Feldblum S, Patel N, Tobin AJ (1991) Two genes encode distinct glutamate decarboxylases. Neuron 7:91–100.

Flores-Herr N, Protti DA, Wassle H (2001) Synaptic currents generating the inhibitory surround of ganglion cells in the mammalian retina. J Neurosci 21:4852–4863.

Forster D, Helmbrecht TO, Mearns DS, Jordan L, Mokayes N, Baier H (2020) Retinotectal circuitry of larval zebrafish is adapted to detection and pursuit of prey. Elife 9.

Forster D, Arnold-Ammer I, Laurell E, Barker AJ, Fernandes AM, Finger-Baier K, Filosa A, Helmbrecht TO, Kolsch Y, Kuhn E, Robles E, Slanchev K, Thiele TR, Baier H, Kubo F (2017) Genetic targeting and anatomical registration of neuronal populations in the zebrafish brain with a new set of BAC transgenic tools. Sci Rep 7:5230.

Gabriel JP, Trivedi CA, Maurer CM, Ryu S, Bollmann JH (2012) Layer-specific targeting of direction-selective neurons in the zebrafish optic tectum. Neuron 76:1147–1160.

Gaetz W, Bloy L, Wang DJ, Port RG, Blaskey L, Levy SE, Roberts TP (2014) GABA estimation in the brains of children on the autism spectrum: measurement precision and regional cortical variation. Neuroimage 86:1–9.

Gahtan E, Tanger P, Baier H (2005) Visual prey capture in larval zebrafish is controlled by identified reticulospinal neurons downstream of the tectum. J Neurosci 25:9294–9303.

Galanopoulou AS (2010) Mutations affecting GABAergic signaling in seizures and epilepsy. Pflugers Arch 460:505–523.

Ganguly K, Schinder AF, Wong ST, Poo M (2001) GABA itself promotes the developmental switch of neuronal GABAergic responses from excitation to inhibition. Cell 105:521–532.

Garyfallidis E, Brett M, Amirbekian B, Rokem A, Van Der Walt S, Descoteaux M, Nimmo-Smith I (2014) Dipy, a library for the analysis of diffusion MRI data. Frontiers in Neuroinformatics 8.

Geschwind DH (2009) Advances in autism. Annual review of medicine 60:367–380.

Glykys J, Dzhala VI, Kuchibhotla KV, Feng G, Kuner T, Augustine G, Bacskai BJ, Staley KJ (2009) Differences in cortical versus subcortical GABAergic signaling: a candidate mechanism of electroclinical uncoupling of neonatal seizures. Neuron 63:657–672.

Grama A, Engert F (2012) Direction selectivity in the larval zebrafish tectum is mediated by asymmetric inhibition. Front Neural Circuits 6:59.

Grone BP, Maruska KP (2016) Three Distinct Glutamate Decarboxylase Genes in Vertebrates. Sci Rep 6:30507.

Grone BP, Qu T, Baraban SC (2017) Behavioral Comorbidities and Drug Treatments in a Zebrafish scn1lab Model of Dravet Syndrome. eNeuro 4.

Guizar-Sicairos M, Thurman ST, Fienup JR (2008) Efficient subpixel image registration algorithms. Opt Lett 33:156–158.

Helmbrecht TO, Dal Maschio M, Donovan JC, Koutsouli S, Baier H (2018) Topography of a Visuomotor Transformation. Neuron 100:1429–1445 e1424.

Higashijima S-I, Mandel G, Fetcho JR (2004) Distribution of prospective glutamatergic, glycinergic, and GABAergic neurons in embryonic and larval zebrafish. J Comp Neurol 480:1–18.

Hildebrand DGC et al. (2017) Whole-brain serial-section electron microscopy in larval zebrafish. Nature 545:345–349.

Hindriks R, Adhikari MH, Murayama Y, Ganzetti M, Mantini D, Logothetis NK, Deco G (2016) Can sliding-window correlations reveal dynamic functional connectivity in resting-state fMRI? Neuroimage 127:242–256.

Holmes GL, Ben-Ari Y (2001) The neurobiology and consequences of epilepsy in the developing brain. Pediatr Res 49:320–325.

Hunter PR, Lowe AS, Thompson ID, Meyer MP (2013) Emergent properties of the optic tectum revealed by population analysis of direction and orientation selectivity. J Neurosci 33:13940–13945.

Isaacson JS, Strowbridge BW (1998) Olfactory reciprocal synapses: dendritic signaling in the CNS. Neuron 20:749–761.

James N, Liu X, Bell A (2016) A fluorescence in situ hybridization (FISH) protocol for stickleback tissue. Evol Ecol Res 17:603–617.

Kaufmann A, Mickoleit M, Weber M, Huisken J (2012) Multilayer mounting enables long-term imaging of zebrafish development in a light sheet microscope. Development 139:3242.

Kim J, Namchuk M, Bugawan T, Fu Q, Jaffe M, Shi Y, Aanstoot HJ, Turck CW, Erlich H, Lennon V, Baekkeskov S (1994) Higher autoantibody levels and recognition of a linear NH2-terminal epitope in the autoantigen GAD65, distinguish stiff-man syndrome from insulin-dependent diabetes mellitus. The Journal of experimental medicine 180:595–606.

Kimmel CB, Ballard WW, Kimmel SR, Ullmann B, Schilling TF (1995) Stages of embryonic development of the zebrafish. Dev Dyn 203:253–310.

Kinoshita M, Ito E (2006) Roles of periventricular neurons in retinotectal transmission in the optic tectum. Prog Neurobiol 79:112–121.

Kinoshita M, Ueda R, Kojima S, Sato K, Watanabe M, Urano A, Ito E (2002) Multiple-site optical recording for characterization of functional synaptic organization of the optic tectum of rainbow trout. Eur J Neurosci 16:868–876.

Koyama M, Pujala A (2018) Mutual inhibition of lateral inhibition: a network motif for an elementary computation in the brain. Curr Opin Neurobiol 49:69–74.

Kramer A, Wu Y, Baier H, Kubo F (2019) Neuronal Architecture of a Visual Center that Processes Optic Flow. Neuron 103:118–132 e117.

Kriegstein AR, Owens DF (2001) GABA may act as a self-limiting trophic factor at developing synapses. Sci STKE 2001:pe1.

Kunst M, Laurell E, Mokayes N, Kramer A, Kubo F, Fernandes AM, Forster D, Dal Maschio M, Baier H (2019a) A Cellular-Resolution Atlas of the Larval Zebrafish Brain. Neuron 103:21–38 e25.

Kunst M, Laurell E, Mokayes N, Kramer A, Kubo F, Fernandes AM, Förster D, Dal Maschio M, Baier H (2019b) A Cellular-Resolution Atlas of the Larval Zebrafish Brain. Neuron 103:21–38.e25.

Lai F, Fagernes Cathrine E, Nilsson Goran E, Jutfelt F (2016) Expression of genes involved in brain GABAergic neurotransmission in three-spined stickleback exposed to near-future CO_2_. Conserv Physiol 4:cow068.

Lai F, Fagernes Cathrine E, Jutfelt F, Nilsson Goran E (2017) Erratum: Expression of genes involved in brain GABAergic neurotransmission in three-spined stickleback exposed to near-future CO_2_. Conserv Physiol 5:cox004.

Lee S, Zhou ZJ (2006) The synaptic mechanism of direction selectivity in distal processes of starburst amacrine cells. Neuron 51:787–799.

Legay F, Pelhate S, Tappaz ML (1986) Phylogenesis of brain glutamic acid decarboxylase from vertebrates: immunochemical studies. J Neurochem 46:1478–1486.

Levy LM, Dalakas MC, Floeter MK (1999) The stiff-person syndrome: an autoimmune disorder affecting neurotransmission of gamma-aminobutyric acid. Ann Intern Med 131:522–530.

Liu J, Baraban SC (2019) Network Properties Revealed during Multi-Scale Calcium Imaging of Seizure Activity in Zebrafish. eNeuro 6.

Liu Y, Dale S, Ball R, VanLeuven AJ, Sornborger A, Lauderdale JD, Kner P (2019a) Imaging neural events in zebrafish larvae with linear structured illumination light sheet fluorescence microscopy. Neurophotonics: SPIE.

Liu Y, Dale S, Ball R, VanLeuven AJ, Sornborger A, Lauderdale JD, Kner P (2019b) Imaging neural events in zebrafish larvae with linear structured illumination light sheet fluorescence microscopy. Neurophotonics 6:015009.

Lynex CN, Carr IM, Leek JP, Achuthan R, Mitchell S, Maher ER, Woods CG, Bonthon DT, Markham AF (2004) Homozygosity for a missense mutation in the 67 kDa isoform of glutamate decarboxylase in a family with autosomal recessive spastic cerebral palsy: parallels with Stiff-Person Syndrome and other movement disorders. BMC Neurol 4:20.

Manent JB, Demarque M, Jorquera I, Pellegrino C, Ben-Ari Y, Aniksztejn L, Represa A (2005) A noncanonical release of GABA and glutamate modulates neuronal migration. J Neurosci 25:4755–4765.

Mann HB, Whitney DR (1947) On a Test of Whether one of Two Random Variables is Stochastically Larger than the Other. The Annals of Mathematical Statistics 18:50–60, 11.

Marachlian E, Avitan L, Goodhill GJ, Sumbre G (2018) Principles of Functional Circuit Connectivity: Insights From Spontaneous Activity in the Zebrafish Optic Tectum. Front Neural Circuits 12:46.

Marquart GD, Tabor KM, Horstick EJ, Brown M, Geoca AK, Polys NF, Nogare DD, Burgess HA (2017) High-precision registration between zebrafish brain atlases using symmetric diffeomorphic normalization. Gigascience 6:1–15.

Miyata S, Kakizaki T, Fujihara K, Obinata H, Hirano T, Nakai J, Tanaka M, Itohara S, Watanabe M, Tanaka KF, Abe M, Sakimura K, Yanagawa Y (2021) Global knockdown of glutamate decarboxylase 67 elicits emotional abnormality in mice. Mol Brain 14:5.

Mullins M (1995) Genetic nomenclature guide. Zebrafish. Trends Genet:31–32.

Muto A, Ohkura M, Abe G, Nakai J, Kawakami K (2013) Real-time visualization of neuronal activity during perception. Curr Biol 23:307–311.

Naumann EA, Fitzgerald JE, Dunn TW, Rihel J, Sompolinsky H, Engert F (2016) From Whole-Brain Data to Functional Circuit Models: The Zebrafish Optomotor Response. Cell 167:947–960 e920.

Nevin LM, Robles E, Baier H, Scott EK (2010) Focusing on optic tectum circuitry through the lens of genetics. BMC Biol 8:126.

Niell CM, Smith SJ (2005) Functional imaging reveals rapid development of visual response properties in the zebrafish tectum. Neuron 45:941–951.

Nikolaou N, Lowe AS, Walker AS, Abbas F, Hunter PR, Thompson ID, Meyer MP (2012) Parametric functional maps of visual inputs to the tectum. Neuron 76:317–324.

O’Connor MJ, Beebe LL, Deodato D, Ball RE, Page AT, VanLeuven AJ, Harris KT, Park S, Hariharan V, Lauderdale JD, Dore TM (2019) Bypassing Glutamic Acid Decarboxylase 1 (Gad1) Induced Craniofacial Defects with a Photoactivatable Translation Blocker Morpholino. ACS Chem Neurosci 10:266–278.

Oh WJ, Westmoreland JJ, Summers R, Condie BG (2010) Cleft palate is caused by CNS dysfunction in Gad1 and Viaat knockout mice. PloS one 5:e9758.

Patel TP, Man K, Firestein BL, Meaney DF (2015) Automated quantification of neuronal networks and single-cell calcium dynamics using calcium imaging. J Neurosci Methods 243:26–38.

Pietri T, Romano SA, Perez-Schuster V, Boulanger-Weill J, Candat V, Sumbre G (2017) The Emergence of the Spatial Structure of Tectal Spontaneous Activity Is Independent of Visual Inputs. Cell Rep 19:939–948.

Pitrone PG, Schindelin J, Stuyvenberg L, Preibisch S, Weber M, Eliceiri KW, Huisken J, Tomancak P (2013) OpenSPIM: an open-access light-sheet microscopy platform. Nat Meth 10:598–599.

Popova E (2015) GABAergic neurotransmission and retinal ganglion cell function. J Comp Physiol A Neuroethol Sens Neural Behav Physiol 201:261–283.

Puts NA, Harris AD, Crocetti D, Nettles C, Singer HS, Tommerdahl M, Edden RA, Mostofsky SH (2015) Reduced GABAergic inhibition and abnormal sensory symptoms in children with Tourette syndrome. J Neurophysiol 114:808–817.

Radmanesh M, Jalili M, Kozlowska K (2020) Activation of Functional Brain Networks in Children With Psychogenic Non-epileptic Seizures. Front Hum Neurosci 14:339.

Ramdya P, Engert F (2008) Emergence of binocular functional properties in a monocular neural circuit. Nat Neurosci 11:1083–1090.

Randlett O, Wee CL, Naumann EA, Nnaemeka O, Schoppik D, Fitzgerald JE, Portugues R, Lacoste AM, Riegler C, Engert F, Schier AF (2015) Whole-brain activity mapping onto a zebrafish brain atlas. Nat Methods 12:1039–1046.

Represa A, Ben-Ari Y (2005) Trophic actions of GABA on neuronal development. Trends Neurosci 28:278–283.

Robertson CE, Ratai EM, Kanwisher N (2016) Reduced GABAergic Action in the Autistic Brain. Curr Biol 26:80–85.

Robles E, Smith SJ, Baier H (2011) Characterization of genetically targeted neuron types in the zebrafish optic tectum. Front Neural Circuits 5:1.

Robles E, Filosa A, Baier H (2013) Precise lamination of retinal axons generates multiple parallel input pathways in the tectum. J Neurosci 33:5027–5039.

Robles E, Laurell E, Baier H (2014) The retinal projectome reveals brain-area-specific visual representations generated by ganglion cell diversity. Curr Biol 24:2085–2096.

Ronneberger O, Liu K, Rath M, Ruebeta D, Mueller T, Skibbe H, Drayer B, Schmidt T, Filippi A, Nitschke R, Brox T, Burkhardt H, Driever W (2012) ViBE-Z: a framework for 3D virtual colocalization analysis in zebrafish larval brains. Nat Methods 9:735–742.

Ross MK, Filipov NM (2006) Determination of atrazine and its metabolites in mouse urine and plasma by LC-MS analysis. Anal Biochem 351:161–173.

Satou C, Kimura Y, Hirata H, Suster ML, Kawakami K, Higashijima S (2013) Transgenic tools to characterize neuronal properties of discrete populations of zebrafish neurons. Development 140:3927–3931.

Schoppa NE, Urban NN (2003) Dendritic processing within olfactory bulb circuits. Trends Neurosci 26:501–506.

Scott EK, Baier H (2009) The cellular architecture of the larval zebrafish tectum, as revealed by gal4 enhancer trap lines. Front Neural Circuits 3:13.

Scott EK, Mason L, Arrenberg AB, Ziv L, Gosse NJ, Xiao T, Chi NC, Asakawa K, Kawakami K, Baier H (2007) Targeting neural circuitry in zebrafish using GAL4 enhancer trapping. Nat Methods 4:323–326.

Solimena M, Folli F, Aparisi R, Pozza G, De Camilli P (1990) Autoantibodies to GABA-ergic neurons and pancreatic beta cells in stiff-man syndrome. The New England journal of medicine 322:1555–1560.

Stuermer CA (1988) Retinotopic organization of the developing retinotectal projection in the zebrafish embryo. J Neurosci 8:4513–4530.

Swanson OK, Maffei A (2019) From Hiring to Firing: Activation of Inhibitory Neurons and Their Recruitment in Behavior. Frontiers in Molecular Neuroscience 12.

Tao L, Lauderdale JD, Sornborger AT (2011) Mapping Functional Connectivity between Neuronal Ensembles with Larval Zebrafish Transgenic for a Ratiometric Calcium Indicator. Front Neural Circuits 5:2.

Tepper JM, Wilson CJ, Koos T (2008) Feedforward and feedback inhibition in neostriatal GABAergic spiny neurons. Brain Res Rev 58:272–281.

Thisse C, Thisse B (2008) High-resolution in situ hybridization to whole-mount zebrafish embryos. Nature protocols 3:59–69.

Thompson AW, Vanwalleghem GC, Heap LA, Scott EK (2016) Functional Profiles of Visual-, Auditory-, and Water Flow-Responsive Neurons in the Zebrafish Tectum. Curr Biol 26:743–754.

Tremblay R, Lee S, Rudy B (2016) GABAergic Interneurons in the Neocortex: From Cellular Properties to Circuits. Neuron 91:260–292.

VanLeuven AJ, Park S, Menke DB, Lauderdale JD (2018) A PAGE screening approach for identifying CRISPR-Cas9-induced mutations in zebrafish. Biotechniques 64:275–278.

Vanwalleghem GC, Ahrens MB, Scott EK (2018) Integrative whole-brain neuroscience in larval zebrafish. Curr Opin Neurobiol 50:136–145.

Vucinic D, Cohen LB, Kosmidis EK (2006) Interglomerular center-surround inhibition shapes odorant-evoked input to the mouse olfactory bulb in vivo. J Neurophysiol 95:1881–1887.

Wang Y, Xu C, Xu Z, Ji C, Liang J, Wang Y, Chen B, Wu X, Gao F, Wang S, Guo Y, Li X, Luo J, Duan S, Chen Z (2017) Depolarized GABAergic Signaling in Subicular Microcircuits Mediates Generalized Seizure in Temporal Lobe Epilepsy. Neuron 95:92–105 e105.

Weber M, Mickoleit M, Huisken J (2014) Multilayer Mounting for Long-term Light Sheet Microscopy of Zebrafish. Journal of Visualized Experiments : JoVE:51119.

Westerfield M (1993) The zebrafish book: a guide for the laboratory use of zebrafish (Brachydanio rerio). Eugene, OR: M. Westerfield.

Westerfield M, ed (2007) The zebrafish book: A guide for the laboratory use of zebrafish (Danio rerio), 5 Edition. Eugene, OR: University of Oregon Press.

Wu C, Sun D (2015) GABA receptors in brain development, function, and injury. Metab Brain Dis 30:367–379.

Wu Y, Dal Maschio M, Kubo F, Baier H (2020) An Optical Illusion Pinpoints an Essential Circuit Node for Global Motion Processing. Neuron.

Xiao T, Baier H (2007) Lamina-specific axonal projections in the zebrafish tectum require the type IV collagen Dragnet. Nat Neurosci 10:1529–1537.

Xiao T, Roeser T, Staub W, Baier H (2005) A GFP-based genetic screen reveals mutations that disrupt the architecture of the zebrafish retinotectal projection. Development 132:2955–2967.

Yokoi M, Mori K, Nakanishi S (1995) Refinement of odor molecule tuning by dendrodendritic synaptic inhibition in the olfactory bulb. Proc Natl Acad Sci U S A 92:3371–3375.

Yoon JW, Yoon CS, Lim HW, Huang QQ, Kang Y, Pyun KH, Hirasawa K, Sherwin RS, Jun HS (1999) Control of autoimmune diabetes in NOD mice by GAD expression or suppression in beta cells. Science 284:1183–1187.

Yu M, Xi Y, Pollack J, Debiais-Thibaud M, Macdonald RB, Ekker M (2011) Activity of dlx5a/dlx6a regulatory elements during zebrafish GABAergic neuron development. Int J Dev Neurosci 29:681–691.

Zalesky A, Fornito A, Cocchi L, Gollo LL, Breakspear M (2014) Time-resolved resting-state brain networks. Proceedings of the National Academy of Sciences 111:10341–10346.

Zhang M, Liu Y, Wang SZ, Zhong W, Liu BH, Tao HW (2011) Functional elimination of excitatory feedforward inputs underlies developmental refinement of visual receptive fields in zebrafish. J Neurosci 31:5460–5469.

