## Supplemental Information for "Decreased GABA levels during development result in increased connectivity in the larval zebrafish tectum"

*Generation of zebrafish mutant for gad1a and gad1b*

sgRNA design: The sequences for CRISPR target sites in the *gad1a* and *gad1b* genes were designed using the ZiFiT Targeter program website (http://zifit.partners.org/ZiFiT/ChoiceMenu.aspx) (Hwang et al., 2013a; Hwang et al., 2013b). The resultant target sequences for each gene used in this study can be found in Supplemental Table 1. These target sequences were evaluated for the probability of causing off-target effects using the ZiFiT program. These sequences were confirmed via nucleotide BLAST searches against the zebrafish genome to contain at least three or more mismatches at any other loci thereby limiting the possibility of off-target effects.

sgRNA, Cas9 and RNP construction: The sgRNA for targeting *gad1b* (*ga2303* allele) was made from a PCR-amplified template as described in (Nakayama et al., 2013) and transcribed using the MEGAshortscriptTM T7 Kit (AM1354, Thermo Fischer Scientific, Inc., Waltham, MA) following the manufacturer protocol. The Cas9 mRNA was transcribed from an optimized expression vector as described in (Jao et al., 2013) using the mMessage mMACHINE Sp6 Kit (AM1340, Thermo Fischer Scientific, Inc., Waltham, MA) following the manufacturer protocol. Both the sgRNA and the Cas9 mRNA were purified by LiCl precipitation and re-dissolved in DEPC-treated water.

The *gad1a (ga2404)* allele was made using a ribonucleoprotein (RNP) complex (Burger et al., 2016). To synthesize this sgRNA, we ordered gene-specific crRNA targeting *gad1a* exon5 as well as trRNA which can be used with any crRNA. Both of these RNA oligonucleotides were purchased from Integrated DNA Technologies (IDT, Skokie, IL) following the online ordering protocol and upon arrival were resuspended to a concentration of 100 µM in the provided Nuclease Free Duplex Buffer. Purified Cas9 nuclease was also purchased from IDT (1074181). Immediately before performing embryonic microinjections, we made a working dilution of Cas9 nuclease in Cas9 working buffer (20 mM HEPES, 150 mM KCl, pH 7.5). The sgRNA duplex was made by combining equimolar amounts (3.4 µL each) of crRNA and trRNA plus 3.2 µL of Nuclease Free Duplex Buffer, incubating at 95°C for 5 minutes, and cooling to room temperature on the benchtop. The RNP complex is assembled by combining 1 µL each of the sgRNA duplex and the Cas9 protein dilution, 0.5 µL DEPC-treated water and 2.5 µL 0.4M KCl with phenol red and incubating at 37°C for 10 minutes.

*Microinjections*

sgRNA for gad1b exon4 plus Cas9 mRNA were co-injected into 1-2 cell stage zebrafish embryos in an approximately 1 nL solution that contained 100 ng/µL of sgRNA and 600 ng/µL of Cas9 mRNA. The RNP complex for gad1a was injected into 1-2 cell stage zebrafish embryos in an approximately 1 nL solution such that the concentration of the sgRNA duplex is 234 pg (156 pg trRNA, 78 pg crRNA) and the concentration of the Cas9 enzyme is 736 pg. In all cases, only animals that exhibited normal development post-injection were grown to adulthood as potential F0 founders.

*Identification and description of gad1a$^{-/-}$ and gad1b$^{-/-}$ mutant zebrafish*

Potential founders that were injected with CRISPR-Cas9 material were crossed with wild-type fish and the F1 progeny were genotyped via PCR and PAGE analyses (VanLeuven et al., 2018). Animals that showed evidence of a mutation based upon this initial screening were then sequenced via Sanger sequencing to determine the nature of the mutation (Fig. S1). Animals with the same mutation were housed together and bred to identify fish with homozygous mutations for each allele. F2 generations were sequence-verified before being established as secured homozygous mutant lines (Supplemental Figure 1). Progeny of the F2 generation were grown to adulthood and were used for all behavioral and molecular aspects of this study.

*Casper, tg(elavl3:gcamp5G) and gad1b-/-, tg(elavl3:gcamp5G) generation and genotyping*

*nacre*$^{w2/w2}$, e*lavl3*:gCaMP5G were crossed to crystal fish (*nacre*$^{w2/w2}$, *alb*$^{b4/b4}$, *roy*$^{a9/a9}$) and the progeny were grown to adults. Progeny were then in-crossed to generate casper and crystal fish expressing e*lavl3*:gCaMP5G. However, crystal fish were never obtained, therefore only casper or nacre fish were used. Crispr generated *gad1b-/-* fish, as previously mentioned, were crossed to *nacre*$^{w2/w2}$, tg(e*lavl3*:gCaMP5G) fish to generate *gad1b* +/-, *elavl3*:gCaMP5G tg/o fish. The progenies were grown to adulthood and in-crossed to generate *gad1b -/-*, *elavl3*:gCaMP5G fish. In this process, it was realized that gad1b and nacre reside on the same chromosome, so no *gad1b-/-*, *nacre* $^{w2/w2}$ fish were ever obtained. Mutants were identified by screening through PCR and PAGE as previously mentioned.

*Fish rearing for calcium imaging*
Larvae from a cross of casper (*nacre*$^{w2/w2,}$ *roy*$^{a9/a9}$ ) fish crossed to casper (*nacre*$^{w2/w2,}$ *roy*$^{a9/a9}$ ), e*lavl3*:gCaMP5G tg/o were used for the wildtype fish and wildtype fish treated with 15mM PTZ groups. For the mutants, larvae were used from a cross of *gad1b -/-*, e*lavl3*:gCaMP5G fish crossed to *gad1b -/-*. As the *gad1b-/- fish* are not in the nacre background, they were treated with 0.003% PTU in egg water starting at 18 hpf. The solution was changed once daily until 5dpf.

*In Situ Hybridization probe construction*
Probes were constructed for *gad1b* by PCR amplification from whole-embryo cDNA using primers listed in supplemental table 1. Probes for *gad1a* and *gad2* were constructed from gBlock Gene Fragments whose sequences are in Supplemental table 1) DIG-labeled riboprobes for all genes were synthesized with a DIG Labeling Kit (Roche 11175025910) per the manufacturer's protocol. Antisense probes were generated using the T7 promoter and sense probes were generated using the Sp6 promoter.

*Colorimetric whole mount in situ hybridization*
Whole-mount in situ hybridizations were performed as previously described on 3dpf and 5dpf WIK zebrafish treated with 0.003% PTU (Thisse and Thisse, 2008). Hybridizations were done overnight in hybridization buffer in a 68°C water bath. Color development was performed using NBT/BCIP (Roche) substrate.

*Colorimetric sectioned in situ hybridization*
5dpf wik zebrafish fixed in 4%pfa were prepared for cryosection and sliced into 10 μm sections. The in situ was performed as a modified version of what is described in Thisse and Thisse. An antigen retrieval step was added as described in (James et al., 2016).

*Image Preprocessing*

To address any potential artifacts resulting from the larvae's motion during the light-sheet imaging experiments, we first applied the "subpixel image registration" procedure (Guizar-Sicairos et al., 2008) to correct for movement of the sample during imaging. In this process, the first frame of the video was considered the target image. The subsequent frames of the video were aligned with the first frame by optimizing the cross-correlation between the registered subsequent frames and the first frame.

Afterward, we proceeded to identify and correct potential change points resulting from fluctuations in the environmental light conditions. To accomplish this, we extracted the background signal by computing the average signal from the pixels situated at the four square corners (7.8 μm × 7.8 μm) of the images. The detection of temporal change points was achieved using the "Pelt" function from the Python package "ruptures" version 1.0.2 (Truong et al., 2020) utilizing default arguments for model fitting and setting the argument "pen" to 10 for prediction. Subsequently, for each identified temporal change point, denoted as $t$, we individually computed the intensity change level for each pixel. This was accomplished by calculating the difference between the median fluorescence intensity of the pixel during the time interval from $t-2$ seconds to $t$ and the time interval from $t$ to $t+2$ seconds. Then we corrected the artifacts caused by the environmental light conditions for each pixel by subtracting the estimated intensity change level attributed to each change point. Finally, we applied K-nearest neighbor (KNN) smoothing method to denoise the signals and remove the potential outliers of the signal. The KNN smoothing also enhancd the robustness of our framework against the estimation error caused during the change point detection procedure. The number of neighbors is set as 7.

Next, we normalized the signal of each pixel by calculating fluorescence intensity change (ΔF/F). This normalization procedure also helped eliminate the decaying effect of the fluorescence. Whole frame average intensities were computed for each image time series. The first 100 frames (4.4 seconds) were omitted from this computation because the LSM excitation light initially activates neural activity in the larvae. Fluorescence intensity changes (ΔF/F) were then calculated using the sliding window method described in (Patel et al., 2015; Liu and Baraban, 2019). This method involves finding the median value in the window interval before each data point ($F_{t0}$), subtracting the mean ($F_{\mu}$) of the data points below the median from the original data point($F_{t0-\Delta t}$), then normalizing by dividing this result by the same mean value. This process is described by the following equation:

$$\Delta F/F = \frac{F_{t0} - F_{\mu}}{F_{\mu}}$$

*Image Analysis*

We performed a Fourier Transform on the one-dimensional fluorescence traces to get the frequency spectra. Then we calculated the absolute value of the real part of the frequency spectrum and normalized it to the maximum value of the frequency spectrum. This analysis was performed for three fish each from three different groups. One group was the fish without any chemical treatment or genetic mutation, here referred to as control. The second group was the fish which were treated with PTZ, here referred to as PTZ-treated. The third group was the genetically mutated fish, here referred to as gad1b$^{-/-}$ mutant (O'Connor et al., 2019).

In order to compare across the changes in neural activity within the optic tectum of the zebrafish larva, we register our image time-series to the zebrafish brain atlas (Kunst et al., 2019) The masks of regions of interests are downloaded from the brain atlas website (https://fishatlas.neuro.mpg.de/).

To determine the position of the imaged plane in 3D, we conducted a search among the atlas images, with the objective of maximizing the cross-correlation with the average image obtained from our collected video.

The brain atlas used in this process was segmented into multiple regions of interest (ROIs). Treating the average brain image from our collected video as the target image, we applied a non-rigid image registration method to align the atlas image with our average brain image. This allowed us to accurately map the positions of the ROIs in our data. The non-rigid image registration is achieved with the “Symmetric Diffeomorphic Registration” method from the python package “dipy” version 1.3.0. (Garyfallidis et al., 2014). Once the ROIs in our dataset were segmented, we proceeded to extract the signal from each ROI. This was accomplished by calculating the average intensity of the image pixels within each ROI.

Subsequently, we proceeded to explore both static and dynamic functional connectivity among the identified ROIs using the extracted signals.
For static functional connectivity, we computed the Pearson correlation directly between the extracted signals from each pair of ROIs. This analysis provided insights into the overall interregional connectivity during the experiments.
To uncover the dynamic nature of the functional interactions among the ROIs, we employed a sliding-window framework, inspired by the work of (Zalesky et al., 2014; Hindriks et al., 2016). This framework allowed us to investigate the dynamic functional connectivity, capturing the variations in interregional connectivity over time. Specifically, we used a tapered window of length 20 s. We slid the window in time at a temporal resolution of 3s over the 5-min interval to get a continuous series of snapshots of the ROI signals. For each snapshot, we calculated a correlation matrix for the ROIs using Pearson correlation. Finally, we obtained a series of correlation matrices (regions × regions × windows). We considered two regions to be connected at any snapshot if the corresponding correlation was larger than 0.6. The frequency of connections between the two sides of brains and within each side of brain were summarized (Fig. 5).

*Code Accessibility*

The code is available here:

[***https://github.com/Knerlab/neural-activity-analysis***](https://github.com/Knerlab/neural-activity-analysis)

### Supplemental Tables

#### Supplemental Table 1: Screening and sequencing primers.

| | Forward (5'-3') | Reverse (5'-3') |
|---|---|---|
| ***gad1a* screening** | CATTAGCATTGACTTGACCGAG | AGCAGGAACTGCATGGTGTA |
| ***gad1a* sequencing** | ACTCAGCGATGCAATGTCAG | TGTGCATGGTCTTCATCACC |
| ***gad1b* screening** | CCCGTGTTGTAATGATGCAG | GTGAAGCGCTCATTGTTGTC |
| ***gad1b* sequencing** | TCCCAGTAAAACTCCAACCG | GCGAACAGGTTGGAGAAATC |

#### Supplemental Table 2: Mean normalized concentration of GABA in adult brains by genotype and comparisons of the mean by one-way ANOVA.

| | WT | $gad1b^{ga2303\ +/-}$ | $gad1b^{ga2303\ -/-}$ | WT and Het ($p < 0.05$) | WT and -/- ($p < 0.05$) | Het and -/- ($p < 0.05$) |
|---|---|---|---|---|---|---|
| GABA | 987 ng/mg | 743.3 ng/mg | 644.6 ng/mg | Yes: $p = 0.0324$ | Yes: $p = 0.0023$ | No |

#### Supplemental Table 3: Mean normalized concentration of neurotransmitters in adult brains by genotype and comparisons of the mean by one-way ANOVA.

| | WT | $gad1b^{ga2303\ +/-}$ | $gad1b^{ga2303\ -/-}$ | WT:Het ($p < 0.05$) | WT:-/- ($p < 0.05$) | Het:-/- ($p < 0.05$) |
|---|---|---|---|---|---|---|
| Serotonin (5-HT) | 0.3025 ng/mg | 0.261 ng/mg | 0.3166 ng/mg | No | No | No |
| Serotonin Metabolite (5-HIAA) | 0.2186 ng/mg | 0.2254 ng/mg | 0.1912 ng/mg | No | No | No |
| Dopamine (DA) | 0.1536 ng/mg | 0.205 ng/mg | 0.1575 ng/mg | No | No | No |
| Glutamate | 1,423 ng/mg | 1,046 ng/mg | 1,252 ng/mg | Yes $p = 0.0022$ | No | No |
| Glutamine | 1,342 ng/mg | 1,138 ng/mg | 1,155 ng/mg | No | No | No |
| Norepinephrine (NE) | 1.051 ng/mg | 1.007 ng/mg | 1.113 ng/mg | No | No | No |
| Norepinephrine Metabolite (MHPG) | 26.81 ng/mg | 31.34 ng/mg | 24.56 ng/mg | No | No | No |

**Supplemental Table 4: Mean normalized concentration of neurotransmitters in 7 dpf larvae by genotype and comparisons of the mean by one-way ANOVA.**

| | WT | *gad1b*$^{ga2303}$ +/- | *gad1b*$^{ga2303}$ -/- | *gad1a*$^{ga2404}$ -/- | WT:Het ($p < 0.05$) | WT:-/- ($p < 0.05$) | Het:-/- ($p < 0.05$) |
|---|---|---|---|---|---|---|---|
| Serotonin (5-HT) | 0.073 ng/mg | 0.035 ng/mg | 0.015 ng/mg | n/a | No | No | No |
| Serotonin Metabolite (5-HIAA) | 1.485 ng/mg | 1.625 ng/mg | 0.8693 ng/mg | n/a | No | No | No |
| GABA | 60.59 ng/mg | 25.17 ng/mg | 56.61 ng/mg | 0.6367 ng/mg | No | No | No |
| Glutamate | 324.1 ng/mg | 477.1 ng/mg | 357.8 ng/mg | 294.8 ng/mg | No | No | No |
| Glutamine | 716.8 ng/mg | 724.8 ng/mg | 3,259 ng/mg | 724.8 ng/mg | No | No | No |
| Norepinephrine (NE) | 0.5743 ng/mg | 0.1233 ng/mg | 0.414 ng/mg | n/a | No | No | No |
| Norepinephrine Metabolite (MHPG) | 1,081 ng/mg | 1,107 ng/mg | 614.5 ng/mg | n/a | No | No | No |
| Dopamine (DA) | n/a | n/a | n/a | n/a | n/a | n/a | n/a |

### Supplemental Figures

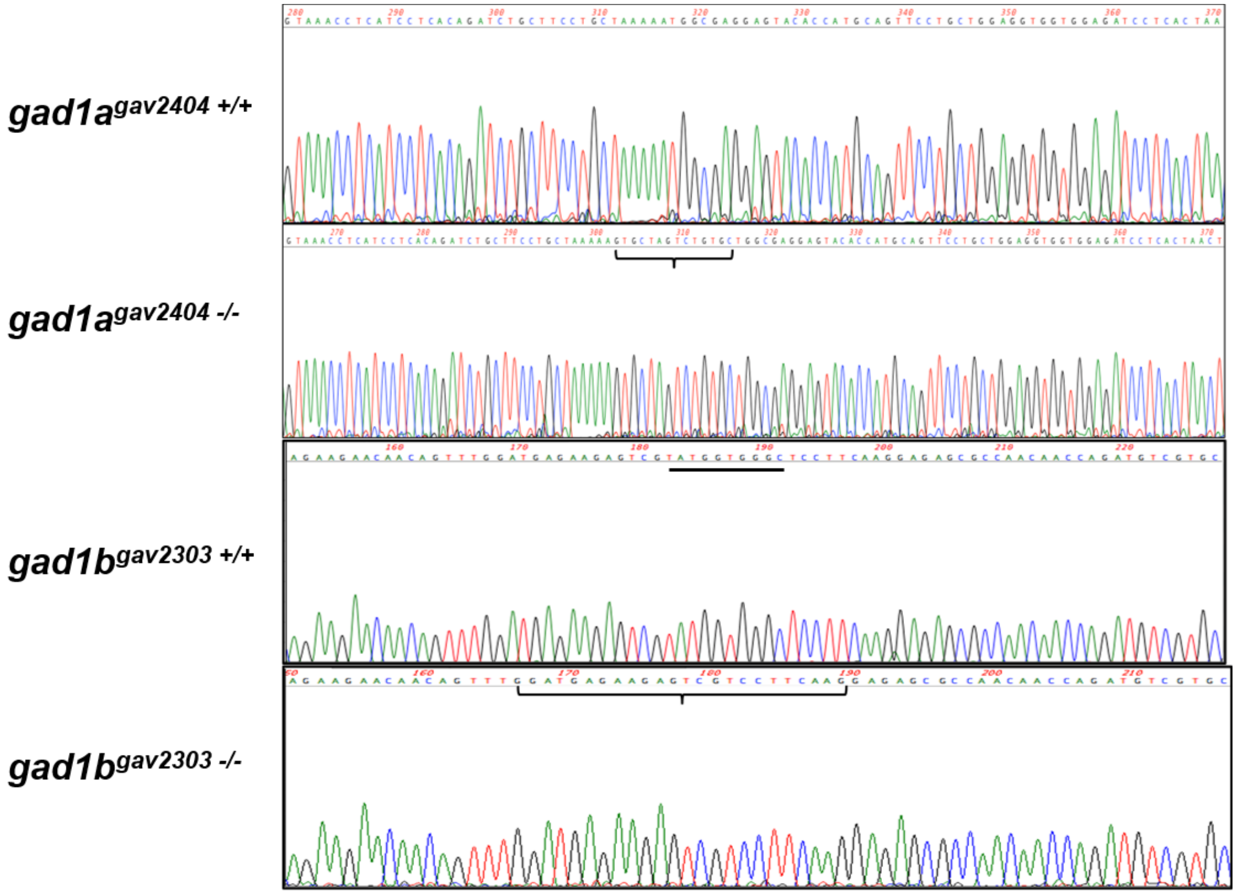


**Figure S1: Sequence for *gad1a*[*ga2404*] and *gad1b*[*ga2303*] alleles.** Sequence verification of homozygous mutations in *gad1a* and *gad1b* that we generated. The bracket in *gad1a* -/- represents the 14 bp insertion that is not present in wild-type. The black line in *gad1b* +/+ indicates the 10 bp that are deleted. The bracket in *gad1b* -/- shows the sequence with the 10 bp deletion.

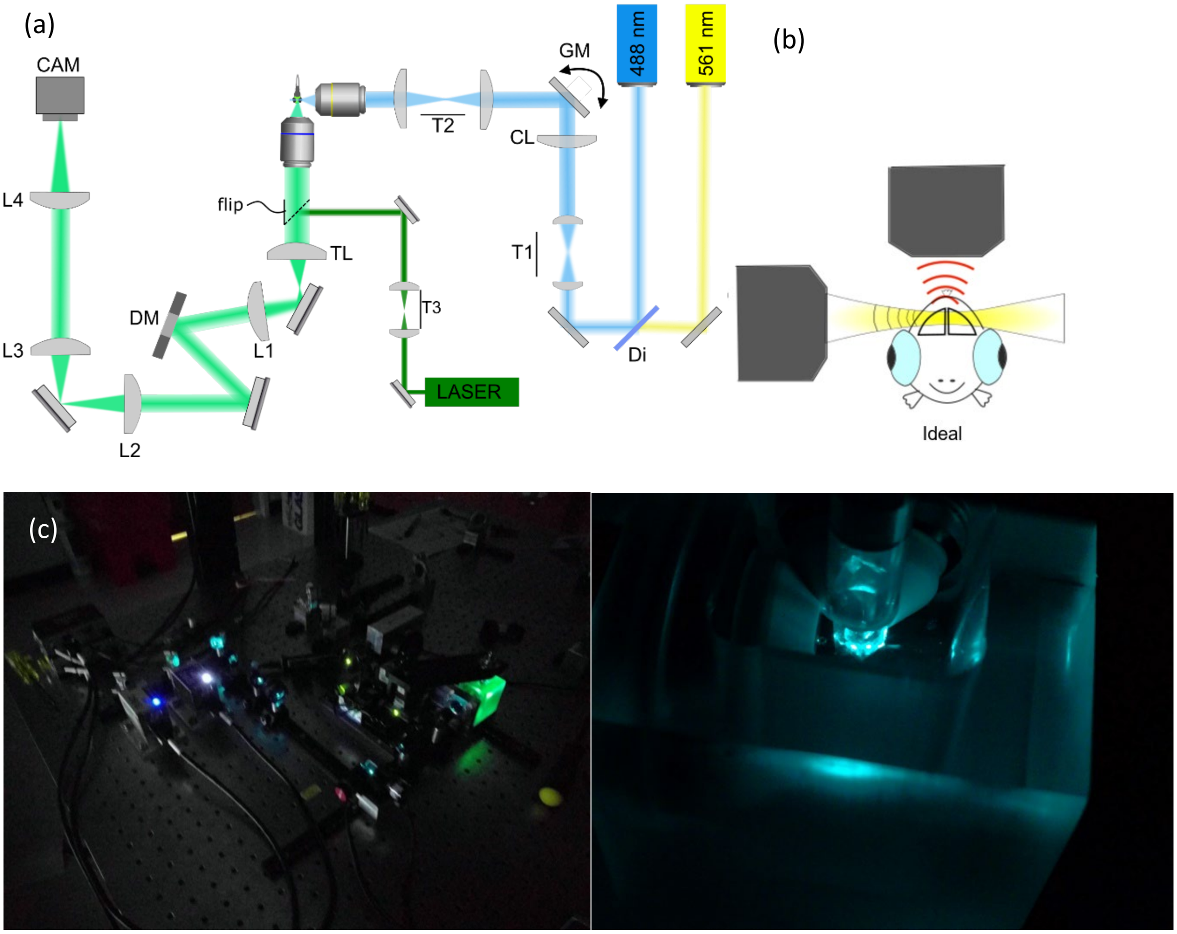


**Figure S2. Schematic of the optical setup.** (a) We use a 10x 0.3NA and a 20X 0.5NA Olympus watering dipping objective lens for illumination and detection, respectively (UMPLFLN10XW, Olympus UMPLFLN20XW). A cylindrical lens (f=50mm) in the illumination path forms an elliptical beam at the back-pupil plane of the 10X objective lens. This creates a thin static Gaussian light sheet with thickness (full width half maximum) of 6.6 µm. The system is capable of two-color imaging with a 488nm laser (Coherent OBIS LX 50 mW) and a 561nm laser (Coherent OBIS LS 50 mW). The final magnification of the system is either 26.67 or 33.3. The 26.67 magnification gives an effective pixel size of 244 nm and a Field of view of 500 × 500 $\mu m^2$. The 33.3 magnification gives an effective pixel size of 195 nm and a Field of view of 399 × 399 $\mu m^2$. The camera (Hamamatsu ORCA flash 4.0 V2) is operated at 33-50 frames per second for single channel imaging. Di: dichroic mirror (Thorlabs DMLP505T). T1: 2x magnification lens pair (Thorlabs AC127-025-A-ML and AC127-050-A-ML). CL: cylindrical lens (Thorlabs ACY254-050-A). T2: 5/3 demagnification lens pair (Thorlabs AC127-050-A-ML and AC127-030-A-ML). TL: tube lens (Olympus U-TLU 180 mm efl). L1: achromatic lens, 80 mm or 100 mm efl (Thorlabs AC508-080-A-ML or AC508-100-A-ML). DM: Deformable Mirror (Boston Micromachines 140-actuator Multi-3.5-DM). With the L1=100mm, a plain mirror was used in place of the DM. L2: achromatic lens 200 mm efl (Thorlabs AC508-200-A-ML). L3: achromatic lens, 300 efl (Thorlabs ACT508-300-A-ML). L4: achromatic lens 300 mm efl (Thorlabs AC508-200-A-ML). GV: two-axis galvo mirror (Thorlabs GVSM002). A green He-Ne laser and T3: magnification lens pair (25mm and 200mm efl) is used for alignment. A multibandpass filter (Semrock FF01-514/605/730-25) is placed in front of the camera. (b) Schematic of larva orientation relative to excitation and imaging objectives. (c) Photos of the microscope and imaging chamber.

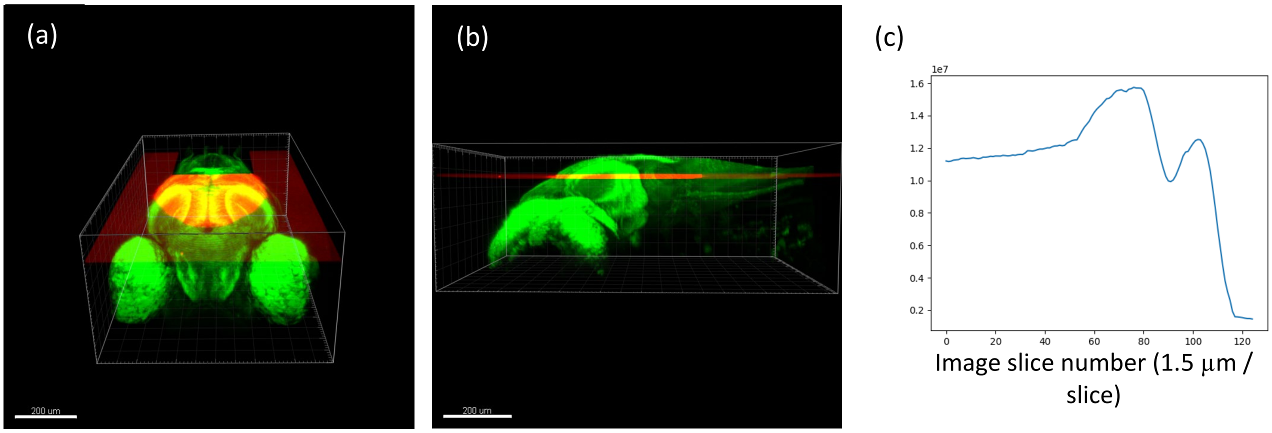


**Figure S3. Target imaging plane.** The target imaging plane is in the middle of the optic tectum. The location of the plane is determined by a cross-correlation of the 2D image (red) with a 3D image of the brain (green). The 2D image was cross-correlated with each slice of the 3D image as shown in (c). The imaging plane was determined to be located at 75 to 81 μm from the dorsal surface – slice 118 in (c). Scale bar in (a) and (b) is 200 μm.

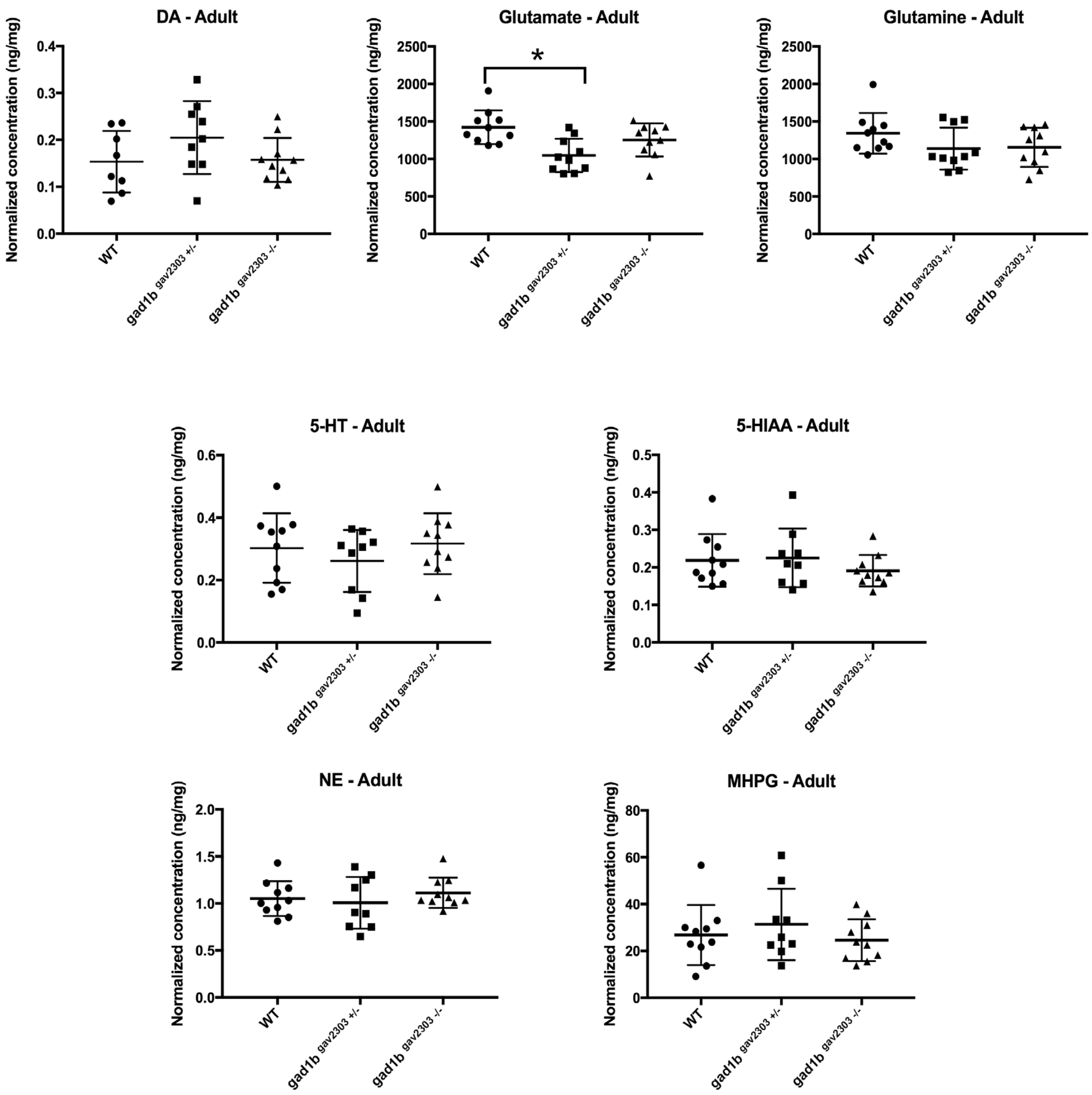


**Figure S4: HPLC-ECD showing the levels other neurotransmitters tested in WT, *gad1b*$^{ga2303\ +/-}$ and *gad1b*$^{ga2303\ -/-}$ adult zebrafish brains.** Normalized concentrations of 7 other neurotransmitters tested in the three genotypes. Abbreviations: 5-HT = serotonin; 5-HIAA = a serotonin metabolite; DA = dopamine; NE = norepinephrine; MHPG = a norepinephrine metabolite. There is no statistically significant change in any of these levels of neurotransmitter across these genotypes except for a decrease in glutamate between WT and *gad1b* +/- (noted with an asterisk).

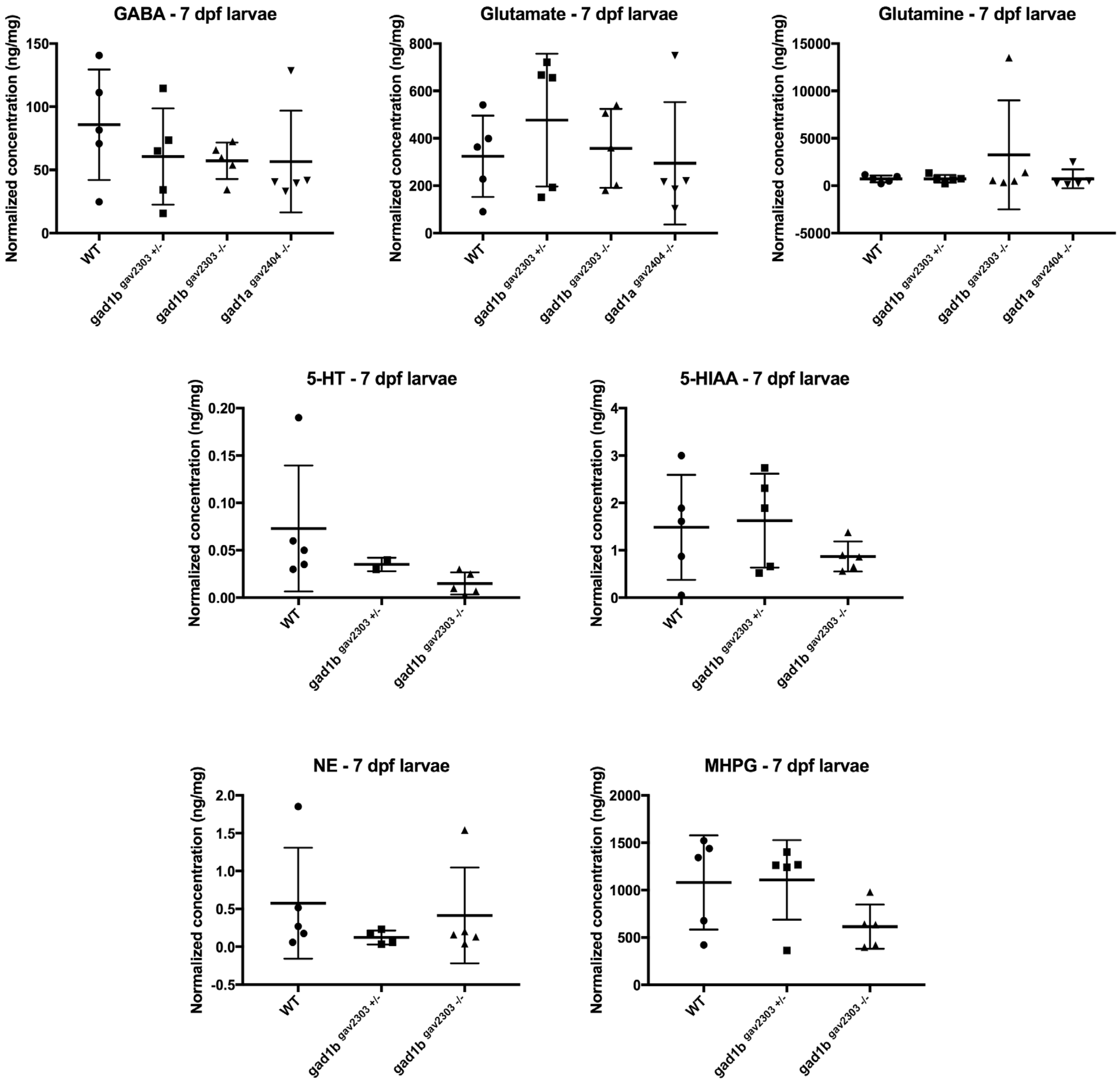


**Figure S5: HPLC-ECD showing the levels other neurotransmitters tested in 7 dpf WT, *gad1b*$^{ga2303\ +/-}$, *gad1b*$^{ga2303\ -/-}$ and *gad1a*$^{ga2404\ -/-}$ larvae.** Normalized concentrations of all neurotransmitters tested in the three genotypes. Abbreviations: 5-HT = serotonin; 5-HIAA = a serotonin metabolite; NE = norepinephrine; MHPG = a norepinephrine metabolite. There is no statistically significant change in any of these levels of neurotransmitter across these genotypes. There is frequently a large amount of variation among these samples.

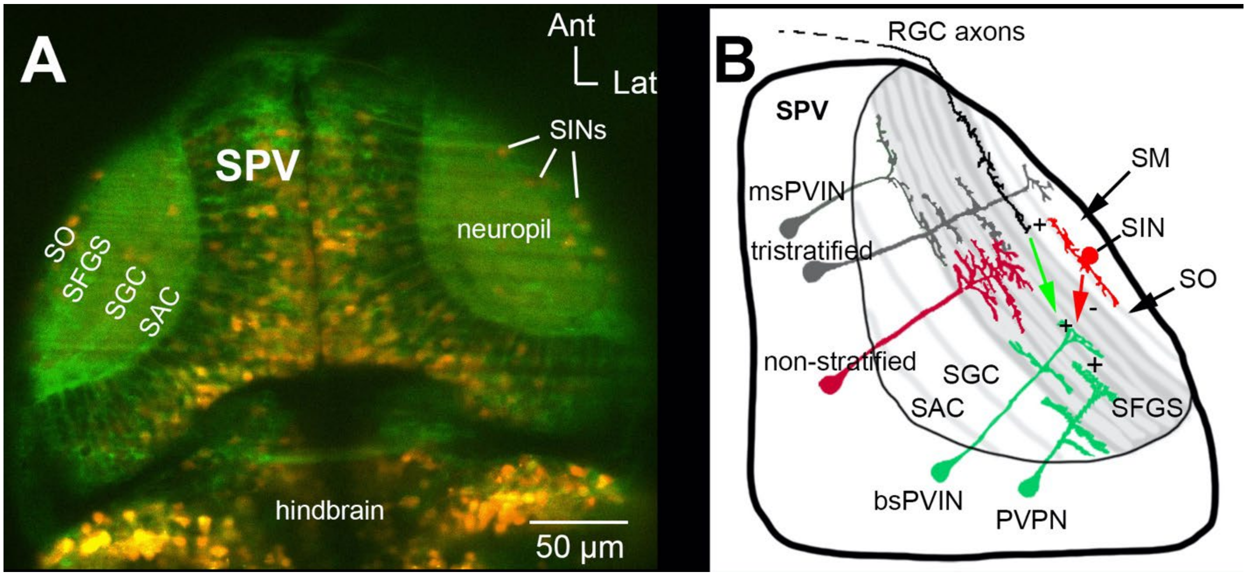


**Figure S6.** (A) Dorsal view of the optic tectum of a 6 dpf larval zebrafish Tg[*elavl3:GCaMP5g*]; TgBAC[*gad1b:loxP-DsRed-loxP-GFP*]. All neurons express GCaMP5g (green). Neurons expressing TgBAC[*gad1b:loxP-DsRed-loxP-GFP*] (Satou et al., 2013) are in red. (B) Cartoon of the tectum showing cell types and neuropil layers. SINs, bsPVINs, and PVPNs form a hypothetical microcircuit (Nevin et al., 2010). stratum opticum (SO); stratum fibrosum et griseum superficiale (SFGS); stratum griseum centrale (SGC); stratum album centrale (SAC); stratum periventriculare (SPV)

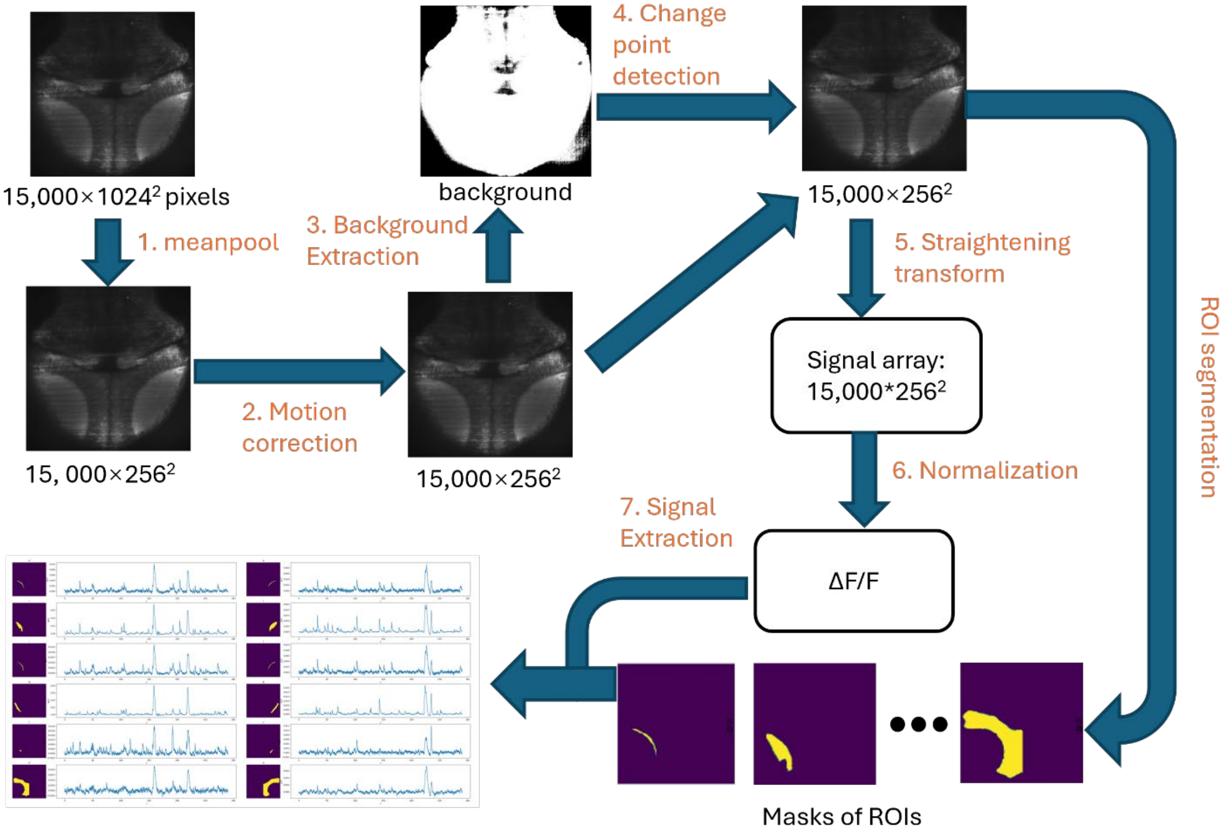


**Figure S7. Workflow for Preprocessing Calcium Imaging and Signal Extraction:** Step 1: Mean pooling of the video to denoise the data. Step 2: Apply motion correction to correct any movement of the Fish during imaging. Step 3: Extract the background signal to detect and correct any disruptions or artifacts introduced by environmental lighting. Step 4: Detect changes in calcium level. Step 5: Straighten images to align with canonical image of the zebrafish CNS. Step 6: Normalize the signal at each pixel. Identify and segment the regions of interest (ROIs). Step 7: Extract the main signal from each identified ROI.

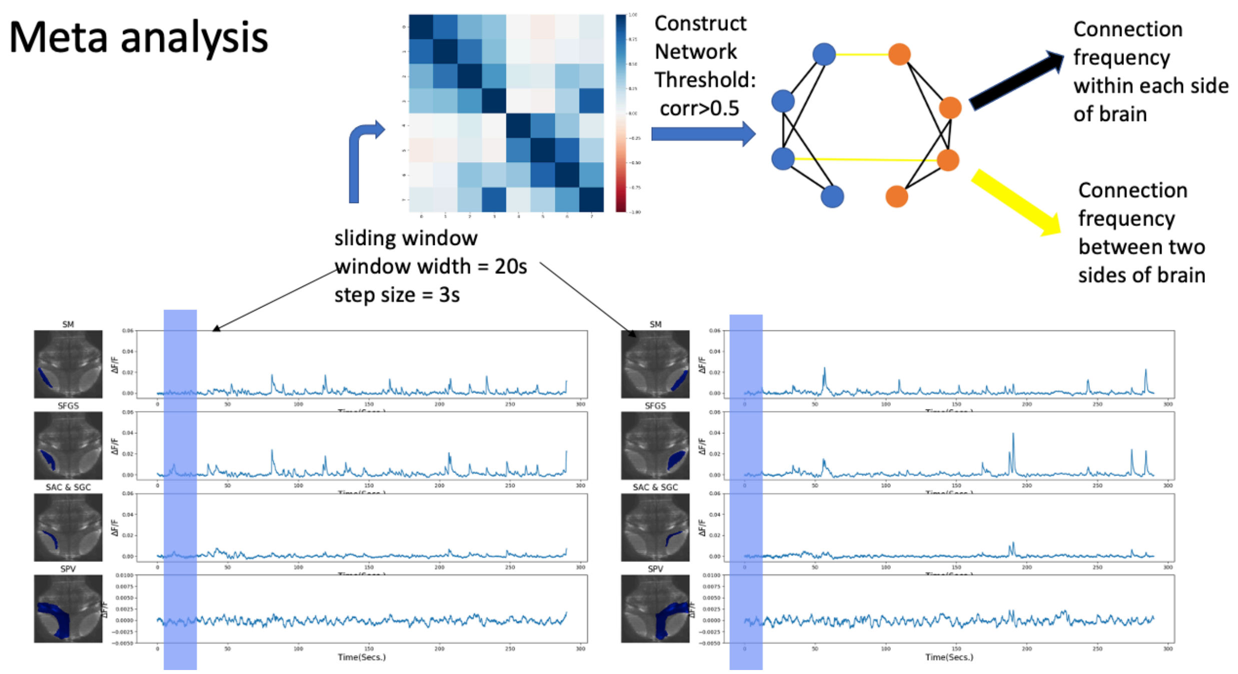


**Figure S8. Meta analysis.** The sliding window technique is applied to assess dynamic functional connectivity across eight regions of interest. At each time point, a Pearson correlation matrix is computed to evaluate the significance of connectivity between each pair of regions. The dynamic network, reflecting these functional connections, is then constructed by applying a 0.6 threshold to the Pearson correlation matrix. The connection frequencies of each imaging sample are used in the down-stream analysis.

### Supplemental References


Ahrens MB, Huang KH, Narayan S, Mensh BD, Engert F (2013a) Two-photon calcium imaging during fictive navigation in virtual environments. Front Neural Circuits 7:104.

Ahrens MB, Orger MB, Robson DN, Li JM, Keller PJ (2013b) Whole-brain functional imaging at cellular resolution using light-sheet microscopy. Nat Methods 10:413-420.

Antinucci P, Hindges R (2016) A crystal-clear zebrafish for in vivo imaging. Sci Rep 6:29490.

Baraban SC, Dinday MT, Hortopan GA (2013) Drug screening in Scn1a zebrafish mutant identifies clemizole as a potential Dravet syndrome treatment. Nat Commun 4:2410.

Baraban SC, Taylor MR, Castro PA, Baier H (2005) Pentylenetetrazole induced changes in zebrafish behavior, neural activity and c-fos expression. Neuroscience 131:759-768.

Bruford EA, Braschi B, Denny P, Jones TEM, Seal RL, Tweedie S (2020) Guidelines for human gene nomenclature. Nat Genet 52:754-758.

Bult CJ, Blake JA, Smith CL, Kadin JA, Richardson JE, Mouse Genome Database G (2019) Mouse Genome Database (MGD) 2019. Nucleic Acids Res 47:D801-D806.

Burger A, Lindsay H, Felker A, Hess C, Anders C, Chiavacci E, Zaugg J, Weber LM, Catena R, Jinek M, Robinson MD, Mosimann C (2016) Maximizing mutagenesis with solubilized CRISPR-Cas9 ribonucleoprotein complexes. Development 143:2025-2037.

Coban A, Filipov NM (2007) Dopaminergic toxicity associated with oral exposure to the herbicide atrazine in juvenile male C57BL/6 mice. J Neurochem 100:1177-1187.

Garyfallidis E, Brett M, Amirbekian B, Rokem A, Van Der Walt S, Descoteaux M, Nimmo-Smith I (2014) Dipy, a library for the analysis of diffusion MRI data. Frontiers in Neuroinformatics 8.
Grone BP, Qu T, Baraban SC (2017) Behavioral Comorbidities and Drug Treatments in a Zebrafish scn1lab Model of Dravet Syndrome. eNeuro 4.
Guizar-Sicairos M, Thurman ST, Fienup JR (2008) Efficient subpixel image registration algorithms. Opt Lett 33:156-158.
Hindriks R, Adhikari MH, Murayama Y, Ganzetti M, Mantini D, Logothetis NK, Deco G (2016) Can sliding-window correlations reveal dynamic functional connectivity in resting-state fMRI? Neuroimage 127:242-256.
Hwang WY, Fu Y, Reyon D, Maeder ML, Kaini P, Sander JD, Joung JK, Peterson RT, Yeh J-RJ (2013a) Heritable and Precise Zebrafish Genome Editing Using a CRISPR-Cas System. PloS one 8:e68708.
Hwang WY, Fu Y, Reyon D, Maeder ML, Tsai SQ, Sander JD, Peterson RT, Yeh JRJ, Joung JK (2013b) Efficient genome editing in zebrafish using a CRISPR-Cas system. Nature Biotechnology 31:227-229.
James N, Liu X, Bell A (2016) A fluorescence in situ hybridization (FISH) protocol for stickleback tissue. Evol Ecol Res 17:603-617.
Jao L-E, Wente SR, Chen W (2013) Efficient multiplex biallelic zebrafish genome editing using a CRISPR nuclease system. Proceedings of the National Academy of Sciences 110:13904-13909.
Kaufmann A, Mickoleit M, Weber M, Huisken J (2012) Multilayer mounting enables long-term imaging of zebrafish development in a light sheet microscope. Development 139:3242.
Kimmel CB, Ballard WW, Kimmel SR, Ullmann B, Schilling TF (1995) Stages of embryonic development of the zebrafish. Dev Dyn 203:253-310.
Kunst M, Laurell E, Mokayes N, Kramer A, Kubo F, Fernandes AM, Förster D, Dal Maschio M, Baier H (2019) A Cellular-Resolution Atlas of the Larval Zebrafish Brain. Neuron 103:21-38.e25.
Liu J, Baraban SC (2019) Network Properties Revealed during Multi-Scale Calcium Imaging of Seizure Activity in Zebrafish. eNeuro 6.
Liu Y, Dale S, Ball R, VanLeuven AJ, Sornborger A, Lauderdale JD, Kner P (2019) Imaging neural events in zebrafish larvae with linear structured illumination light sheet fluorescence microscopy. Neurophotonics 6:015009.
Mann HB, Whitney DR (1947) On a Test of Whether one of Two Random Variables is Stochastically Larger than the Other. The Annals of Mathematical Statistics 18:50-60, 11.
Monge-Acuña AA, Fornaguera-Trías J (2009) A high performance liquid chromatography method with electrochemical detection of gamma-aminobutyric acid, glutamate and glutamine in rat brain homogenates. Journal of Neuroscience Methods 183:176-181.
Mullins M (1995) Genetic nomenclature guide. Zebrafish. Trends Genet:31-32.
Nakayama T, Fish MB, Fisher M, Oomen-Hajagos J, Thomsen GH, Grainger RM (2013) Simple and efficient CRISPR/Cas9-mediated targeted mutagenesis in Xenopus tropicalis. genesis 51:835-843.
Nevin LM, Robles E, Baier H, Scott EK (2010) Focusing on optic tectum circuitry through the lens of genetics. BMC Biol 8:126.
O'Connor MJ, Beebe LL, Deodato D, Ball RE, Page AT, VanLeuven AJ, Harris KT, Park S, Hariharan V, Lauderdale JD, Dore TM (2019) Bypassing Glutamic Acid Decarboxylase 1 (Gad1) Induced Craniofacial Defects with a Photoactivatable Translation Blocker Morpholino. ACS Chem Neurosci 10:266-278.

Patel TP, Man K, Firestein BL, Meaney DF (2015) Automated quantification of neuronal networks and single-cell calcium dynamics using calcium imaging. J Neurosci Methods 243:26-38.
Pitrone PG, Schindelin J, Stuyvenberg L, Preibisch S, Weber M, Eliceiri KW, Huisken J, Tomancak P (2013) OpenSPIM: an open-access light-sheet microscopy platform. Nat Meth 10:598-599.
Ross MK, Filipov NM (2006) Determination of atrazine and its metabolites in mouse urine and plasma by LC-MS analysis. Anal Biochem 351:161-173.
Satou C, Kimura Y, Hirata H, Suster ML, Kawakami K, Higashijima S (2013) Transgenic tools to characterize neuronal properties of discrete populations of zebrafish neurons. Development 140:3927-3931.
Thisse C, Thisse B (2008) High-resolution in situ hybridization to whole-mount zebrafish embryos. Nature protocols 3:59-69.
Truong C, Oudre L, Vayatis N (2020) Selective review of offline change point detection methods. Signal Process 167.
VanLeuven AJ, Park S, Menke DB, Lauderdale JD (2018) A PAGE screening approach for identifying CRISPR-Cas9-induced mutations in zebrafish. Biotechniques 64:275-278.
Weber M, Mickoleit M, Huisken J (2014) Multilayer Mounting for Long-term Light Sheet Microscopy of Zebrafish. Journal of Visualized Experiments : JoVE:51119.
Westerfield M (1993) The zebrafish book : a guide for the laboratory use of zebrafish (Brachydanio rerio). Eugene, OR: M. Westerfield.
Westerfield M, ed (2007) The zebrafish book: A guide for the laboratory use of zebrafish (Danio rerio), 5 Edition. Eugene, OR: University of Oregon Press.
Zalesky A, Fornito A, Cocchi L, Gollo LL, Breakspear M (2014) Time-resolved resting-state brain networks. Proceedings of the National Academy of Sciences 111:10341-10346.
